# 4-Phenylbutyric acid modulates Connexin 43 expression restricting murine-β-coronavirus infectivity and virus-induced demyelination

**DOI:** 10.1101/2024.12.10.627737

**Authors:** Grishma Kasle, Madhav Sharma, Saurav Kumar, Pranati Das, Subhajit Das Sarma, Michael Koval, Anita Mahadevan, Lawrence C. Kenyon, Jayasri Das Sarma

## Abstract

Gap junction intercellular communication, particularly involving Connexin 43 and Connexin 47, plays a critical role in maintaining CNS homeostasis and has been implicated in Multiple Sclerosis (MS) pathology. Thus, warranting further studies in experimental animal models to understand how modulation of Cx43 expression can influence MS pathology. Intracranial infection with murine-β-coronavirus Mouse Hepatitis Virus (MHV-A59) in mice results in acute pathology characterized by high viral titers, glial activation in the brain and chronic neuroinflammatory demyelination, effectively mimicking key pathological hallmarks of MS and serving as a robust model to investigate its viral etiology. MHV-A59 infection leads to a pronounced downregulation of Cx43 during the acute phase, emphasizing its critical role in virus-induced CNS pathology. In this study, we investigated the potential of *in vivo* 4-phenylbutyric acid (4-PBA) administration in modulating Cx43 expression in this MHV-induced model and its consequence on chronic virus-induced demyelination. Our results reveal that 4-PBA treatment reduced acute MHV-A59 infectivity and viral spread in the brain while modulating the glial cell response, mounting host immunity. Treatment with 4-PBA effectively preserved the expression of both Cx43 and Cx47 in infected CNS cells, counteracting their infection-induced downregulation. Furthermore, MHV-A59 infection downregulated the expression of ER-resident thioredoxin family protein (ERp29), a well-known molecular chaperone of Cx43, which was rescued by 4-PBA treatment. We further validated if such downregulation of ERp29 is also evident in MS demyelinating plaques. In human MS patient-derived brain tissue, reduced Cx43 and ERp29 staining was observed in demyelinating plaques. Our studies revealed that 4-PBA treatment not only limits viral replication and spread throughout the brain but also protects the mice against severe chronic neuroinflammatory demyelination. These findings suggest that targeting Cx43 with 4-PBA holds significant therapeutic potential for addressing virus-induced neuroinflammatory demyelination and MS by preserving gap junction intercellular communication.

## 1. Introduction

Multiple Sclerosis (MS) is the most common non-traumatic disabling disease of the central nervous system (CNS) affecting young adults with increasing incidence and prevalence in both developed and developing nations [1, 2]. Although the underlying cause of MS remains uncertain, several genetic and environmental risk factors are suggested to play a central role. A number of studies have identified viruses as triggers for MS[3–7]. However, the exact pathophysiology correlating the trigger of viral infection with onset of demyelination is yet to be clearly demonstrated. In this context, experimental animal models of MS have allowed researchers to explore disease initiation and progression mechanisms as well as test several novel therapeutic approaches [2, 5].

Connexin family gap junction proteins, particularly those expressed by glial cells, have been identified as significant contributors to the pathology of MS[8–13]. Connexins assemble into hexameric hemichannels or connexons, which form gap junctions (GJs) when connexons of juxtaposed cells dock together. This enables direct cell-cell communication through GJ channels that interconnect the cytoplasm of adjacent cells. GJs enable the passive intercellular diffusion of small molecules, including glutamate, glutathione, glucose, adenosine triphosphate (ATP), cyclic adenosine monophosphate (cAMP), inositol 1,4,5-trisphosphate (IP3), and ions (Ca^2+^, Na^+^, K^+^) thereby maintaining metabolic coupling within the glial syncytium [14–19].

Various glial cell types express different connexins. Astrocytes express up to three different connexins (Cx43, Cx30, Cx26), as do oligodendrocytes (Cx32, Cx29, Cx47), microglia (Cx43, Cx32, Cx36) and endothelial cells (Cx37, Cx40, Cx43) [14, 20]. Connexin 43(Cx43) is the most highly expressed connexin in the brain because it is involved in extensive GJ coupling between astrocytes, the most abundant CNS cell type. The coupling partner of astrocytic Cx43 is the Cx47 on the oligodendrocytes. This Cx43-Cx47 mediated astrocyte-oligodendrocyte gap junction coupling is crucial for preserving myelin. In MS and neuromyelitis optica (NMO) cases, there was a preferential loss of Cx43 in actively demyelinating as well as chronic active lesions where heterotypic Cx43-Cx47 gap junctions were extensively lost. Further correlating the loss of Cx43 to disease aggressiveness and distal oligodendrogilopathy in both MS and NMO [9].While alterations in the gap junction proteins Cx43 and Cx47 in MS have been documented, their mechanisms are poorly understood, limiting the ability to assess their potential as therapeutic targets. This warrants further studies in experimental animal models to understand how changes in Cx43 expression can influence MS pathology.

Murine β-coronaviruses (CoV), including certain strains of mouse hepatitis virus (MHV-A59, MHV-2 and MHV-JHM) have been widely used as experimental models to gain insight into the mechanism of demyelination in MS. Intracranial inoculation of the hepato-neurotropic stain of MHV, MHV-A59 in 4-week-old C57BL/6 mice causes acute stage infection and neuroinflammation at day 5 post-infection (p.i) followed by chronic stage (day 30 p.i) inflammatory demyelination with or without axonal loss. This neurodegeneration mimics certain pathological features of the human neurological disease MS, making MHV an important animal model for studying virus-induced demyelination and exploring the etiology and pathogenesis of MS [21]. In addition, since MHV is a β- coronavirus, studies in this model will help elucidate mechanisms whereby coronavirus (CoV) infection induces CNS pathology and host response [22, 23].

MHV-A59 infection in the CNS downregulates the expression of Cx43 in the infected CNS, leading to destabilisation and downregulation of Cx47 expression, mimicking their altered expression as observed in MS [24, 25]. MHV-A59 infection and inflammation-mediated alterations in Cx43 and Cx47 expression and their effect on heterotypic GJIC formation between astrocytes and oligodendrocytes are likely to be associated with the observed chronic demyelination in this model. *In vitro* studies have implicated the involvement of an ER-resident thioredoxin family protein (ERp29) in regulating Cx43 expression and trafficking to the cell surface.[26, 27]. However, it remains to be elucidated if ERp29 expression is altered in the CNS during MHV-A59 infection and if that corelates with expression of Cx43 in the infected CNS.

In the current study, we utilise 4-PBA, a chemical chaperone, in an effort to modulate and maintain steady levels of Cx43 expression *in vivo* in the mouse CNS following MHV-A59 infection and study its effect on chronic stage virus-induced demyelination. 4-PBA is a chemical chaperone and histone deacetylase inhibitor that is FDA-approved for treating urea cycle disorders and is under investigation in cancer, hemoglobinopathies, motor neuron diseases, and cystic fibrosis clinical trials[28, 29].While *in vitro* studies have established the ability of 4-PBA to rescue Cx43 expression and trafficking [18, 22, 23], its ability to improve *in vivo* Cx43 expression is yet to be studied.

Our data reveals that 4-PBA treatment reduced acute-stage MHV-A59 infectivity and spread in CNS and modulated the glial cell response to infection. We demonstrate that overall ERp29 expression is downregulated in the infected CNS at acute-stage, with a marked loss in MHV-A59 infected cells. In contrast, 4-PBA-treated mice showed preservation of ERp29 expression even in infected cells. There was a parallel downregulation in Cx43 and Cx47 expression in infected CNS, which was mitigated by 4-PBA treatment. 4-PBA treatment was able to protect mice against chronic-stage virus-induced demyelination. These results highlight the potential of repurposing 4-PBA to address viral-induced demyelination and GJ dysfunction in MS; clinical application of 4- PBA is facilitated by its FDA approval status.

Previous *in vitro* studies, as well as the current study, have associated the downregulation of ERp29 with reduced Cx43 expression, suggesting that reduced ERp29 may likely be an underlying cause of Cx43 downregulation observed in MS patients. The potential for this hypothesis was validated by studying Cx43 and ERp29 expression in demyelinating plaques from MS-patient-derived brain tissue.

## 2. Materials and Methods

### Ethics statement

All experimental procedures and animal care and use were strictly regulated and reviewed in accordance with good animal ethics approved by the Institutional Animal Care and Use Committee at the Indian Institute of Science Education and Research Kolkata (IISERK/IAEC/AP/2023/118). Experiments were performed following the guidelines of the Committee for Control and Supervision of Experiments on Animals (CCSEA), India.

All the experimental procedures involving studies on brain tissues from human MS patients and normal human control tissues were regulated, reviewed, and approved by the Institutional Ethics Committee of Indian Institute of Science Education And Research (IISER) Kolkata (IISER/IEC/2024/10).

### Isolation of primary mixed glial culture and enrichment of astrocytes

Primary mixed glial cell cultures were established from postnatal day 0 to 1 mouse pups, with minor modifications as previously described[30, 31]. Briefly, following the removal of meningeal covering, brain tissues were homogenised and subjected to enzymatic digestion by incubating in a rocking water bath at 37 °C for 30 min in HBSS containing 300 μg/ml DNase I and 10 mg/ml trypsin. Dissociated cells were triturated with 0.25% of FBS in HBSS and passed through a 70-μm nylon mesh. Cells were washed in HBSS at 300g for 10 min and plated in astrocyte growth medium containing DMEM supplemented with 10% FBS, 1% L-glutamine, and 1% penicillin and streptomycin. After 24 hr, media were changed to remove all nonadherent cells, and cells were allowed to grow to confluency with media change every 3 days thereafter. After 9 to 10 days, once the cultures got confluent, the addition of fresh growth medium was stopped for 10 days to allow differential adhesion of astrocytes and microglia. Following this, the culture flasks were thoroughly agitated in an orbital incubator shaker (200 rpm for 40 min at 37 °C), followed by a quick shaking to remove the loosely adherent microglial cells by using differential adherent properties of astrocytes and microglia. The remaining adherent monolayers were enriched in astrocytes.

### Infection of primary astrocytes with MHV-A59 and 4-PBA treatment

Primary astrocytes were infected with inoculation medium (DMEM containing 1% penicillin-streptomycin and 1% glutamine with 2% FBS) containing MHV-A59 at multiplicities of infection (MOI) of 5 and allowed to adhere for 1 hr at 37°C in a humidified CO_2_ incubator. After 1 hr of incubation, the inoculum was removed and infected cells were maintained in astrocyte-specific medium containing 10% normal FBS (-4PBA) or in astrocyte-specific medium containing 10% FBS supplemented with 10 mM 4-PBA for 24hr. At 24hr p.i, the cell culture medium was removed and the cells were fixed using 4% paraformaldehyde (PFA).

### Immunofluorescence of primary astrocytes

Immunofluorescence studies were performed according to the protocol described previously [24], with minor modifications. Primary astrocytes were plated on etched glass coverslips and fixed with 4% paraformaldehyde (PFA). Permeabilization was done with phosphate-buffered saline (PBS) containing 0.5% Triton X-100 and blocked with PBS containing 0.5% Triton X-100 and 2.5% heat-inactivated goat serum (PBS-GS). The cells were incubated with primary antisera diluted in blocking solution for 1 hr, washed, and then labelled with secondary antisera diluted in blocking solution. Cells were then washed with PBS, mounted with mounting medium containing 4′,6-diamidino-2-phenylindole (DAPI; VectaShield, Vector Laboratories), and visualized using a Zeiss confocal microscope (LSM710). Images were acquired and processed with Zen2010 software (Carl Zeiss).

### Virus, inoculation of mice and experimental design

MHV-free, 4-week-old, C57BL/6 male mice were intracranially inoculated with one-half of the 50% lethal dose (one-half LD_50_) of MHV-A59 (2000 pfu). The mice were monitored daily for weight change, signs and symptoms of disease. Clinical disease severity was graded regularly using the following scale: 0, no disease symptoms; 0.5, ruffled fur, possibly slower movement; 1.0, hunched back position with mild ataxia, possibly slower movement; 1.5, hunched back position with mild ataxia and hind-limb weakness, restricted movement; 2, ataxia, balance problem and/or partial paralysis but still able to move; 2.5, one leg completely paralyzed, motility issue but still able to move around with difficulties; 3, severe hunching/wasting/both hind limb paralysis and mobility is severely compromised; 3.5, severe distress, complete paralysis and moribund; 4, dead[32].

Mice were mock-infected with PBS-BSA and were maintained in parallel. Mice were sacrificed at the peak of inflammation/acute stage (day 5 p.i.) and the peak of demyelination/chronic stage (day 30 p.i.). Brain, Spinal cord and liver tissues were harvested for experimentation.

Mice were divided into two groups: one group (MHV-A59) received intracranial (IC) inoculation with MHV-A59 (50% of the LD_50_ dose: 2000PFU) on day 0. The second group (MHV-A59+4-PBA) was infected with the same dose of MHV-A59 and an intraperitoneal (IP) injection of 4-PBA at 200mg/kg of body weight 0.5 hrs prior to IC inoculation. Additionally, the MHV-A59+4-PBA group received daily 4-PBA injections of 200mg/kg of body weight/day until day 4 for acute stage studies and until day 10 for chronic stage studies (Figure 1 A, B). Vehicle control mice (that received IP PBS injection) were also maintained in parallel.

**Figure 1.**
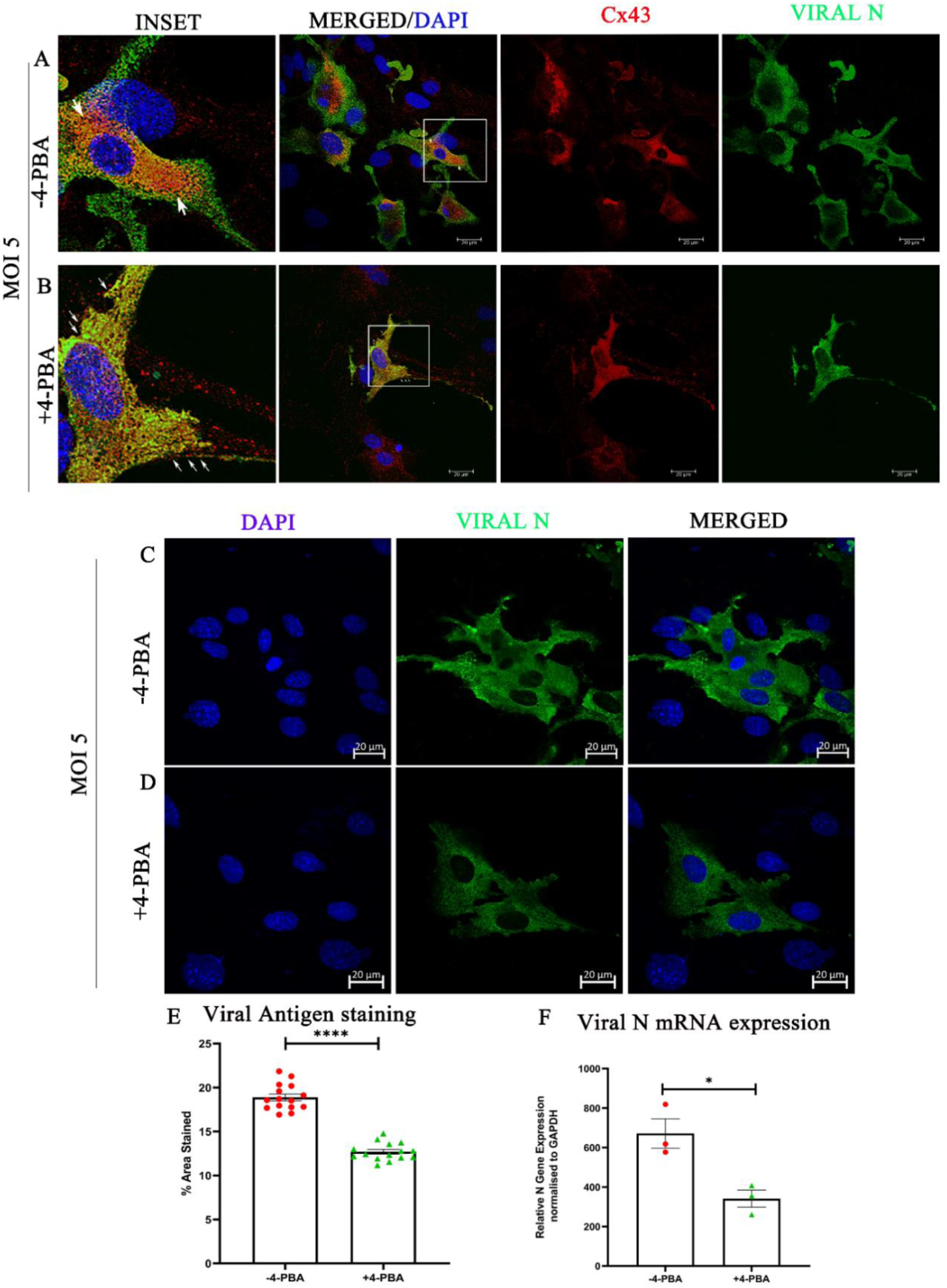
4-PBA treatment improves Cx43 trafficking to the cell surface and reduces MHV-A59 infectivity in mouse primary astrocytes. (A, B) Representative confocal photomicrographs of murine primary astrocytes infected with MHV-A59 at MOI 5 in either the presence(B) or absence(A) of 4-PBA for 24 hp. i. analysed by immunolabeling with anti-Cx43 (red) and anti-nucleocapsid (Viral-N) antibody (green) and mounted in a DAPI-containing mounting medium. The boxed areas are shown alongside as manually zoomed insets. Arrows indicate intracellular retention of Cx43 in (A) and Cx43 puncta in(B) in infected cells. (C-D) Representative confocal photomicrographs of murine primary astrocytes infected with MHV-A59 at MOI 5 in either the presence(D) or absence(C) of 4-PBA for 24 hp. i. analysed by immunolabeling with anti-nucleocapsid (Viral-N) antibody (green) and mounted in a DAPI-containing mounting medium(E) quantification of viral antigen staining in untreated (-4PBA) or 4-PBA treated (+4-PBA) astrocyte cultures. (F) shows the differential viral mRNA expression in untreated or 4-PBA-treated astrocyte cultures infected with MHV-A59. Results were expressed as mean ± SEM (n=3 per group). Asterisks represent statistical significance calculated using unpaired Student’s t-test and Welch correction; p<0.05 was considered significant. *p<0.05, ****p<0.0001.

4-PBA solution was prepared by titrating equimolecular amounts of 4-phenylbutyric acid (Sigma, Madrid, Spain) and sodium hydroxide to pH 7.4. The dose was chosen in reference to a previous dose-response study performed using increasing doses of PBA from 100 to 800 mg/kg to determine the most efficacious dose [33, 34].

### Autopsy tissue and patient characterization

This study was performed on archival autopsied formalin fixed paraffin embedded brain tissue from 1 MS case and 1 age-matched normal control. Tissue sections were obtained from the Human Brain Tissue Repository (Brain Bank), National Institute of Mental Health & Neurosciences, Bangalore 560 029, India for further immunohistochemistry study. The age at autopsy was 21 years for both MS(Female) and Control Case (Male).

### Estimation of viral replication

MHV-A59 and MHV-A59+4-PBA mice were sacrificed at Day 5p.i for estimation of infectious viral particles. Brain and liver tissues were harvested and placed into 1 ml of isotonic saline containing 0.167% gelatin (gel saline). Tissues were weighed and kept frozen at − 80 °C until titered. Tissues were subsequently homogenised, and using the supernatant, viral titers were quantified by standard plaque assay protocol on tight monolayers of L2 cells as described previously [35]using the formula: plaque-forming units (PFUs) = (no. of plaques × dilution factor/ml/gram of tissue) and expressed as log_10_ PFUs/gram of tissue.

### Histopathology and immunohistochemical analysis

Liver, brain, and spinal cord tissues were harvested and embedded in paraffin following transcardial perfusion with PBS. Tissues were post-fixed in 4% paraformaldehyde for 36–48 hrs, following which tissues were processed in increasing concentrations of ethyl alcohol, xylene, and paraffin wax, embedded in paraffin, and sectioned into 5 μm thick transverse sections (liver and spinal cord) and sagittal sections (brain) using Thermo Scientific HM 355 S sectioning system. Liver sections were stained with Hematoxylin and Eosin for histopathologic analysis. Moreover, spinal cords and human brain sections were also stained with Luxol fast blue (LFB) to evaluate for demyelination as described previously[36].

Immunohistochemical staining of the brain (mouse and human) and spinal cord(mouse) tissue serial sections was performed as described previously, using avidin-biotin immunoperoxidase technique (Vector Laboratories) with 3, 3-diaminobenzidine as the substrate[37]. The following primary antibodies were used–anti-Iba-1 (Wako), anti-GFAP (Sigma-Aldrich®, MO, USA),monoclonal antibody directed against the nucleocapsid protein (N) of MHV (monoclonal antibody clone 1-16-1 provided by Julian Leibowitz, Texas A&M University), anti-Cx43(Sigma-Aldrich®, MO, USA) and anti-ERp29(Invitrogen, MA, USA)(Table 2). Control slides from mock-infected and vehicle-control mice were stained in parallel. All the slides were coded and read blindly for subsequent analyses.

### Quantification of histopathological sections

Image analysis was performed using the basic densitometric thresholding application of Fiji (Image J, NIH Image, Scion Image) as described previously [72]. Briefly, image analysis for IHC of Iba1, GFAP and Viral nucleocapsid staining in sections was performed by capturing the images at the highest magnification (4X for brain, 10X for spinal cord) such that the entire section (i.e., scan area) can be visualized within a single frame. The RGB image was deconvoluted into three different colours to separate and subtract the DAB-specific staining from the background Hematoxylin staining. The perimeter of each brain and spinal cord tissue was digitally outlined, and the area was calculated in μm^2^. A threshold value was fixed for each image to ensure that all antibody-marked cells were considered. The amount of Iba1, GFAP and viral nucleocapsid staining was termed as the ‘% area of staining.’

To determine the area of demyelination, LFB-stained spinal cord cross-sections from each mouse were chosen and analysed using Fiji software (Image J, NIH Image, Scion Image) as described previously [72, 73]. Briefly, the total perimeter of the white matter regions in each cross-section was marked and calculated by adding together the dorsal, ventral, and anterior white matter areas in each section. Also, the total area of the demyelinated regions was outlined and collated for each section separately. The percentage of spinal cord demyelination per section per mouse was calculated (5-7 sections from cervical and thoracic regions of the Spinal cord /mouse were examined).

### Protein isolation and western blot analysis

On average, 150mg of brain tissue was collected from euthanized mice following transcardial perfusion with 20ml PBS and flash-frozen in liquid nitrogen. Tissue was then lysed in 1 ml of RIPA buffer (0.1% SDS and 0.1% Triton X-100) with 1X complete mini protease inhibitor cocktail tablets (#11836153001; Roche) and Phosphatase inhibitor cocktail [Sodium Orthovanadate (10Mm), Sodium fluoride(10Mm), Sodium Pyrophosphate(10Mm)]. Brain tissues were homogenized by trituration and sonication. Tissue lysates were centrifuged at 13,500 RPM for 30 min at 4°C, and the supernatant was collected as a whole protein extract. Protein was quantified using Pierce BCA protein assay kit (Thermo Scientific, Rockford, IL, USA). Equal amounts of protein were resolved on SDS-PAGE followed by transfer to polyvinylidene difluoride membranes (Millipore, Bedford, MA) using transfer buffer (25mM Tris, 192mM glycine, and 20% methanol). The membrane was subsequently blocked with 5% non-fat skim milk in TBST (Tris-buffered saline containing 0.1% v/v Tween-20) for 1 hr at room temperature, followed by incubation in respective primary antibodies (Table 2) (in blocking solution) overnight at 4°C. Membranes were then subjected to washes in TBST, followed by incubation with HRP-conjugated secondary IgG. Blots were washed in TBST, and immunoreactive bands were visualised using SuperSignalTM West Pico PLUS Chemiluminescent Substrate, Thermo Fisher Scientific. Densitometric analyses of non-saturated membranes were carried out using a Syngene G: Box chemidoc system and Image J software.

### Gene expression: RNA isolation, reverse transcription, and quantitative polymerase chain reaction

RNA was extracted from the brain or spinal cord tissues (flash-frozen) of MHV-A59, MHV-A59+4-PBA and mock infected mice using the Trizol isolation protocol following transcranial perfusion with DEPC treated PBS. The total RNA concentration was measured using a NanoDrop ND-2000 spectrophotometer. 1μg of RNA was used to prepare cDNA using a High-Capacity cDNA Reverse Transcription Kit (Applied Biosystems).

Quantitative Real-time PCR analysis was performed using iTaq UniverSYBR Green qPCR kit (BioRad) in a BioRad CFX Real-time PCR system (BioRad) under the following conditions: initial denaturation at 95°C for 7min, 40 cycles of 95°C for 10s, 60°C for 30s, melting curve analysis at 60°C for 30s. Reactions were performed in quadruplets. Relative quantitation was performed using the comparative threshold (ΔΔCt) method. mRNA expression levels of target genes in MHV-A59, MHV-A59+4-PBA and Mock infected mice were normalized with GAPDH and expressed as relative fold change compared to their respective mock-infected controls. Primer sequences are enlisted in Table 1.

**Table 1:**
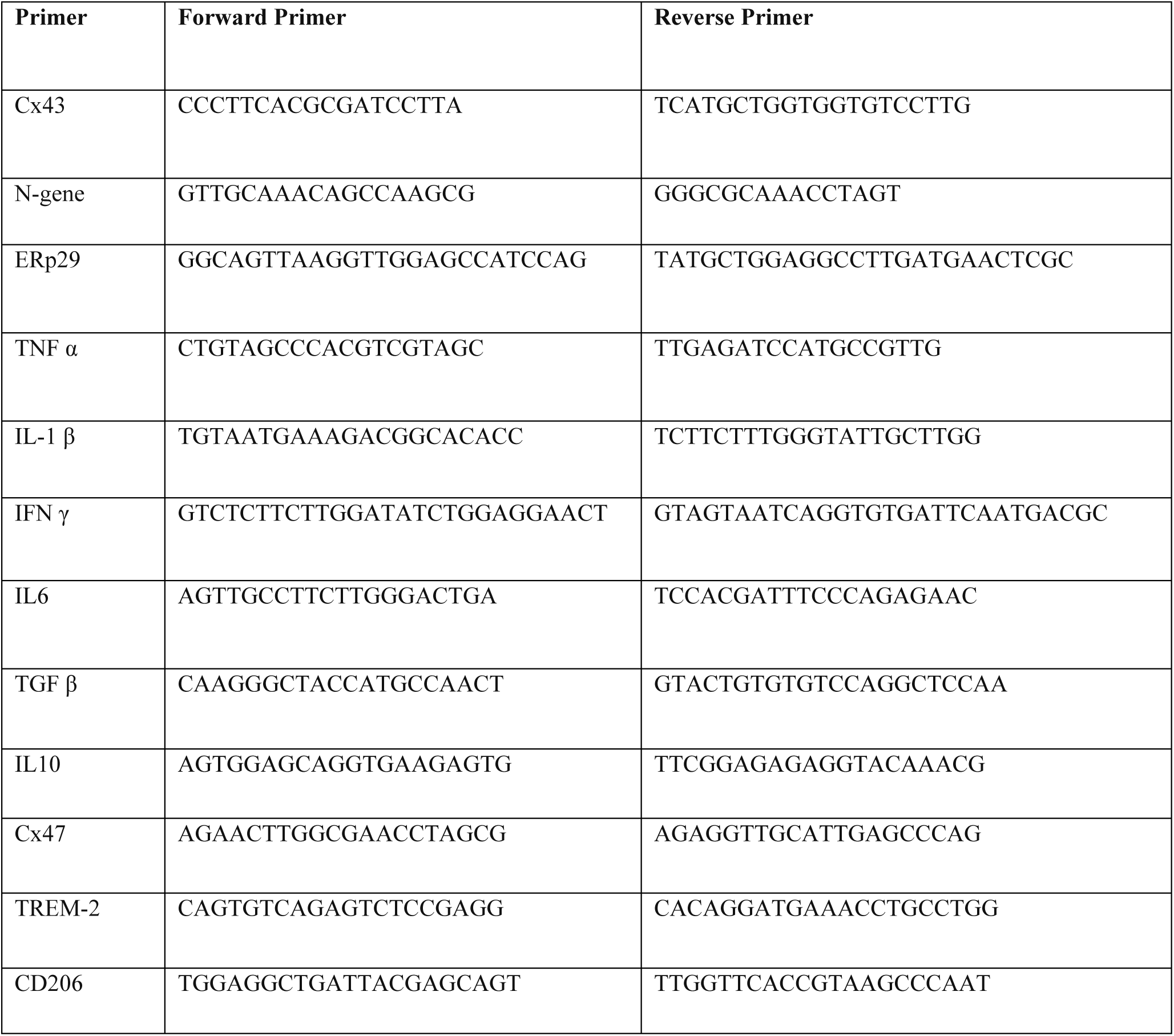
Sequence of Primers used in the study. Primer sequence (5’-3’)

**Table 2:** List of primary antibodies used in the study.

| Sl. No. | Primary antibody | Application | Dilution | Source |
| --- | --- | --- | --- | --- |
| 1 | Anti-Cx43<br>(Polyclonal) | IF<br>WB<br>IHC | 1:1000<br>1:1000<br>1:1000 | Sigma-Aldrich®,<br>MO, USA |
| 2 | Anti-GFAP<br>(Polyclonal) | IF<br>IHC | 1:500<br>1:500 | Sigma-Aldrich®,<br>MO, USA |
| 3 | Anti-Cx47<br>(Polyclonal) | IF<br>WB | 1:500<br>1:1000 | Invitrogen, MA,<br>USA |
| 4 | Anti-Iba1<br>(Polyclonal) | IF<br>IHC | 1:500<br>1:500 | Wako |
| 5 | Anti-Viral<br>Nucleocapsid<br>(monoclonal) | IF<br>IHC | 1:40<br>1:40 | monoclonal<br>antibody clone 1-16-1 provided by<br>Julian Leibowitz,<br>Texas A&M<br>University |
| 6 | Anti-ERp29<br>(Polyclonal) | IF<br>WB<br>IHC | 1:500<br>1:1000<br>1:500 | Invitrogen, MA,<br>USA |

\*\*\*\*\*

### Immunofluorescence microscopy

Brain tissues were harvested and embedded in paraffin following transcranial perfusion with PBS. Tissues were post-fixed in 4% paraformaldehyde for 36–48 hours, following which tissues were processed in increasing concentrations of ethyl alcohol, xylene, and paraffin wax, embedded in paraffin, and sectioned into 5 μm thick Sagittal sections using the Thermo Scientific HM 355 S sectioning system. The tissue sections parallel to those used for corresponding immunohistochemistry were stained. Briefly, the slides were deparaffinized followed by rehydration and antigen unmasking. The slides were then permeabilized with 0.2% Triton X-100 in PBS by shaking incubation at RT for 15 min and blocked using 1% bovine serum albumin (BSA) prepared in 0.2% Triton X-100-1X PBS solution at 37 °C for 1 hr. This was followed by shaking incubation at 4 °C for 16 hr with primary antibodies prepared in blocking solution. Each section on each sample slide was stained for the following separately: i) Anti-Iba1 ii), Anti-GFAP iii) Anti-Iba1 iv) anti-Viral Nucleocapsid v), Anti-Cx43 and anti-Viral Nucleocapsid vi), Anti-Erp29 and anti-Viral Nucleocapsid vii), and Anti-Cx47 and anti-Viral Nucleocapsid. Slides were then washed and incubated for 1 hr at 37 °C with a combination of secondary antibodies AlexaFluor568 and/or AlexaFluor488 prepared in blocking solution. Finally, the slides were washed in PBS and mounted using Vectashield with DAPI.

Images were acquired using a Spinning Disk (Confocal, SORA) Microscope using a 10x objective for Figures 3- 5 and using Leica SP8 confocal platform using an oil immersion 63× objective (NA 1.4) and deconvolved using Leica Lightning software for Figures 7 and 8. The images were captured at a z-interval of 0.2 µm for Figures 7 and 8.

### Statistical analyses

Values were represented as mean values ± standard errors of the mean (SEM). Values were subjected to unpaired student’s t-tests with Welch’s correction or Ordinary One-Way ANOVA with multiple comparison tests (Tukey’s test and the Holm-Sidak test) for calculating the significance of differences between the means. All statistical analyses used GraphPad Prism 8 software (La Jolla, CA). A P-value of < 0.05 was considered statistically significant.

## 3. Results

### 4-PBA treatment improved Cx43 trafficking to the cell surface and reduced MHV-A59 infectivity in mouse primary astrocytes

We previously demonstrated in the astrocytoma-derived cell line, DBT cells, that 4-PBA has the ability to upregulate ERp29, enhance Cx43 trafficking and expression, and reduce the severity of MHV-A59 infectivity and spread [26, 27, 38]. Therefore, we first examined whether 4-PBA treatment had a comparable effect on MHV- A59-infected mouse primary astrocytes *in vitro*.

Primary murine astrocytes were isolated from neonatal mouse pups, infected with MHV-A59 at a multiplicity of infection (MOI) of 5, either untreated (-4-PBA) or treated (+4-PBA) with 4-PBA for 24 hr. Immunofluorescence studies demonstrated that in MHV-A59-infected cells, astrocytes exhibit intracellular retention of Cx43(Figure 1A, insets, retention indicated by arrows). While 4-PBA treatment in MHV-A59 infected astrocytes restored the ability of Cx43 to form puncta at the cell surface (Figure 1B, Insets, puncta indicated by arrows), indicating improved trafficking of Cx43, which was consistent with our previous reports demonstrating that 4-PBA has the ability to promote Cx43 trafficking and expression[26, 27, 38].

Furthermore, the number of infected cells forming syncytia (a morphologic marker of infection) was lower in cells that received 4-PBA treatment (Figure 1D) compared to those that did not receive any treatment (Figure 1C). Quantification of viral antigen staining demonstrated that cells that received 4-PBA treatment showed significantly lower viral antigen staining (Figure 1E). Similarly, viral N gene RNA transcript levels were significantly lower in 4-PBA-treated cultures compared to those that did not receive any treatment (Figure 1F). Thus, 4-PBA reduces MHV-A59 viral infectivity of primary astrocytes *in vitro*. This finding further supported studying the efficacy of 4-PBA treatment against MHV-A59 infection *in vivo*.

### 4-PBA treatment restricts MHV-A59 infection and spread in the brain at the acute stage of infection

The MHV-A59-induced disease in 4-week-old C57BL/6 mice manifests into two temporally distinct stages: acute and chronic. Upon intracranial inoculation with the virus, the disease initially presents with robust viral replication and neuroinflammation accompanied by hepatitis. This acute stage of infection peaks at Day 5 p.i. After day 5 p.i, the viral titer gradually decreases, resolving the neuroinflammation by Day 10 p.i, although the viral RNA persists at low levels. Myelin loss, with or without axonal loss, starts as early as day 7 p.i., and reaches its peak at the chronic phase of infection at 30 days p.i. This neurodegeneration mimics certain pathological features of the human neurological disease MS [21]. Thus, the current study is focused on these two critical time points: the acute stage at Day 5 p.i, and the chronic stage at day 30 p.i.

Mice were divided into two groups: one group (MHV-A59) received intracranial (IC) inoculation with MHV-A59 (50% of the LD50 dose: 2000PFU) on day 0. The second group (MHV-A59+4-PBA) was infected with the same dose of MHV-A59 and received an intraperitoneal (IP) injection of 4-PBA at 200mg/kg of body weight 0.5 hours prior to IC inoculation. Additionally, the MHV-A59+4-PBA group received daily 4-PBA injections of 200mg/kg of body weight/day until day 4 for acute stage studies and until day 10 for chronic stage studies (Figure 2 A, B). Vehicle control mice (that received IP PBS injection) were also maintained in parallel.

**Figure 2.**
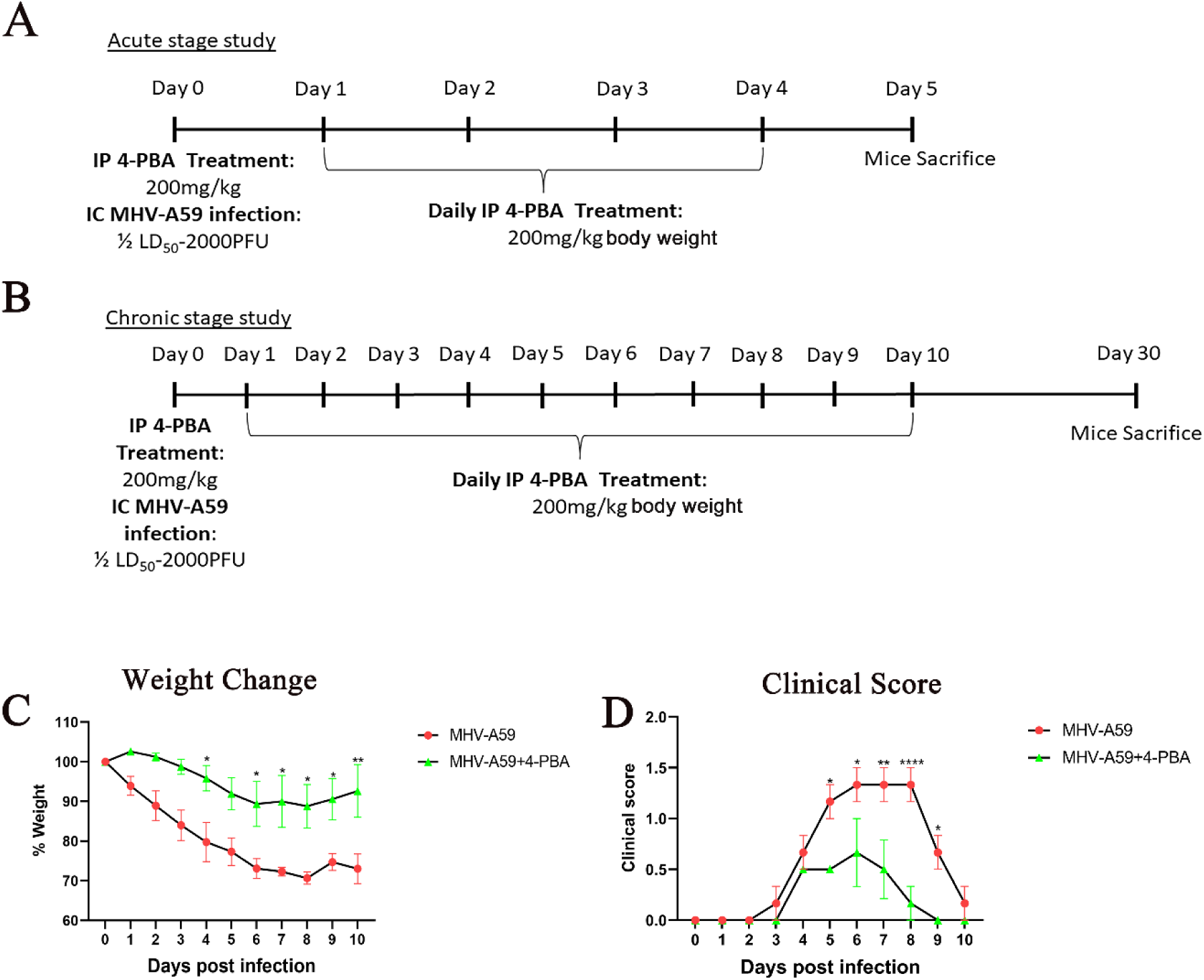
4-PBA treatment improves the weight loss and clinical score in MHV-A59 infected mice. (A,B) Schematic representation of the treatment and infection design. Treated C57BL/6 mice received intraperitoneal injections of 4-PBA(200mg/kg) followed by intracranial inoculation of MHV-A59 (2000 PFU) on day 0 for both acute (A) and Chronic stage studies(B). Day 1 onwards, mice were administered 4-PBA daily up to Day 4 for the acute stage study(A) and up to day 10 for the chronic stage study(B) . Non-injected and mock infected mice were maintained as controls. Mice infected with MHV-A59 that received 4-PBA treatment (MHV-A59+4-PBA) or did not receive any treatment (MHV-A59) were monitored daily for (C) weight change and (D) development of the clinical disease. Clinical scores were assigned by a relative scale of 0–4 as described in Materials and Methods. Results were expressed as mean ± SEM from 2 independent biological experiments (n=6 per group). *Asterisk represents statistical significance calculated using 2-way ANOVA, p<0.05 was considered as significant. *p<0.05, **p<0.01, ****p<0.0001.

The experimental mice were monitored daily for weight change and the development of clinical signs and symptoms. We observed that MHV-A59+4-PBA treated mice showed less than 10% weight loss, while MHV- A59 infected mice lost almost 20% of their original weight by day 4 p.i. (Figure 2 C). MHV-A59 infected mice showed a progressive increase in clinical score starting day 3 p.i., reaching an average score of 1 to 1.5 by day 5- 8 indicated by a hunchback phenotype with the occasional presence of mild ataxia and hind-limb weakness. In contrast, MHV-A59+4-PBA treated mice maintained a lower clinical score with a maximum average of 0.5, indicative of ruffled fur and possibly slower movement (Figure 2 D).

MHV-A59 is a dual hepato-neurotropic virus in which inflammation within the liver is a valuable indicator of disease severity. Acute mild to moderate hepatitis at day 5 p.i. is characteristic of MHV-A59 infection and was observed in infected mice with the presence of multiple necrotizing hepatic lesions spread throughout the liver (Figure 3 A). However, MHV-A59+4-PBA-treated mice showed fewer hepatic lesions, indicative of decreased inflammation in the liver (Figure 3 B). Furthermore, viral titers in the liver of MHV-A59+4-PBA-treated mice were significantly lower compared to MHV-A59-infected mice (Figure 3 C).

**Figure 3.**
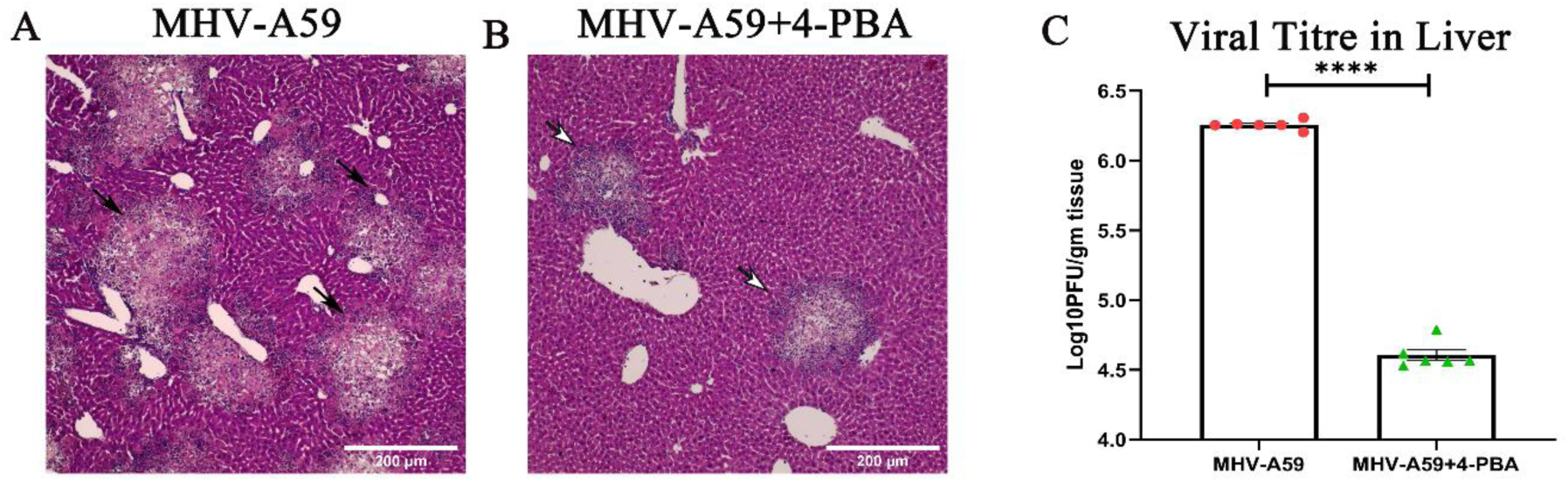
4-PBA treatment reduces acute hepatitis upon MHV-A59 infection. MHV-A59 infection is known to cause acute mild-moderate hepatitis. The reduced weight change and clinical score in MHV-A59+ 4-PBA treated mice prompted investigation of systemic inflammation in the liver. On Day 5p.i MHV-A59 infected(A) and MHV-A59+4-PBA(B) treated mice were subjected to histopathological analyses of liver tissues by H & E staining. Arrows indicate inflamed and necrotic hepatic lesions. (C) Viral titer was determined in liver homogenates at day 5p.i and plotted. Results were expressed as mean ± SEM from 2 independent biological experiments (n=6 per group). *Asterisk represents statistical significance calculated using unpaired Student’s t-test and Welch correction, p<0.05 was considered as significant. ****p<0.0001. Scale bar 200 μm.

To confirm if the reduced disease severity in MHV-A59+4-PBA treated mice correlated with reduced viral spread and infection in the brains of treated mice, we further estimated the viral spread and titers in the brains of experimental mice. Serial paraffin-embedded brain sections from both MHV-A59 infected and MHV-A59+4- PBA treated mice were immunohistochemically stained with anti-viral nucleocapsid antibodies. In correlation with the reduced disease severity, the viral antigen staining was significantly lower in MHV-A59+4-PBA treated mice (1.531% area of staining) as opposed to MHV-A59 (2.932% area of staining) infected mice without 4-PBA treatment (Figure 4 A, B, E). Vehicle control-treated infected mice showed no differential phenotypic or histopathological features compared to MHV-A59 infected mice (Supplementary Figure 1). Immunofluorescence further confirmed that 4-PBA treatment reduced viral spread, including reduced viral antigen staining in the brain (Figure 4 C,D).

**Figure 4.**
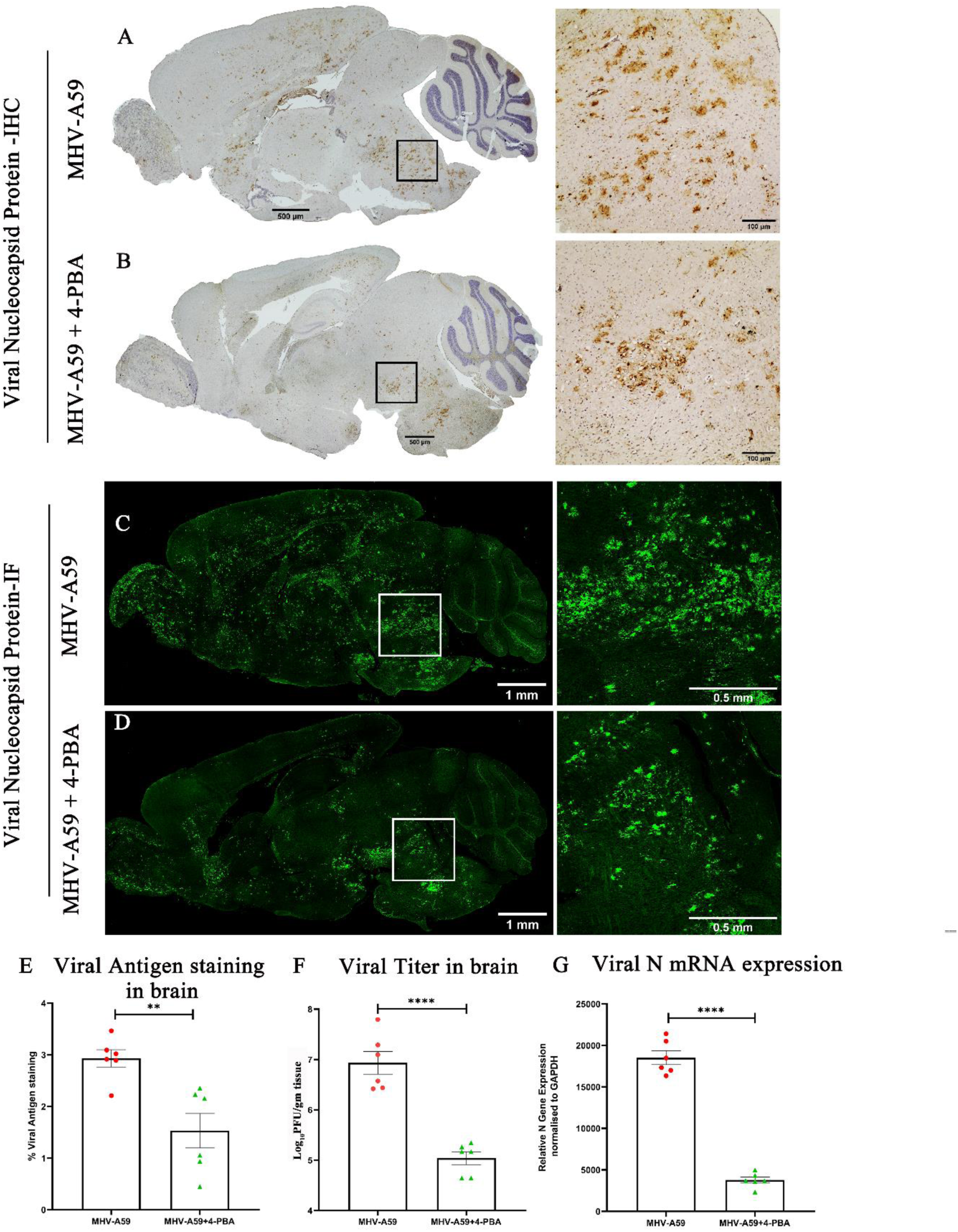
4-PBA treatment restricts in vivo spread and infectivity of MHV-A59 in the mouse brain. (A,B) On day 5p.i , serial mid-sagittal brain sections (5 μm thick) from MHV-A59 (A) and MHV-A59+4-PBA (B) sets of mice were immunohistochemically(IHC) stained with anti-N antibody. The boxed areas are shown at higher magnification alongside the corresponding brain midsagittal sections. (C, D) Photomicrographs of Serial mid-sagittal brain sections (5 μm thick) from MHV-A59(C) and MHV-A59+4-PBA(D) sets of mice stained by immunolabelling with anti-N antibody and visualised using fluorescence. The boxed areas are shown at higher magnification alongside the corresponding brain midsagittal sections (E) Shows quantification of Viral N antigen staining by IHC (A,B) in the brain. (F)Represents comparative viral titres and (G) shows the differential Viral mRNA expression in the brain. Results were expressed as mean ± SEM from 2 independent biological experiments (n=6 per group).*Asterisk represents statistical significance calculated using unpaired Student’s t-test and Welch correction, p<0.05 was considered significant. **p<0.01,****p<0.0001.

To assess if the reduced disease severity and viral antigen staining are correlated with reduced viral replication and infectious viral titers, the brains of MHV-A59 and MHV-A59+4-PBA mice were assayed by plaque assay. Consistent with a reduction in viral antigen spread in response to 4-PBA treatment, brain tissue from MHV- A59+4-PBA treated mice had fewer infectious viral particles than brain tissue from MHV-A59 infected mice (Figure 4 F).

Lastly, we measured the presence of viral mRNA, which correlated with viral antigen and infectious viral particles, where there were fewer viral nucleocapsid transcripts found in MHV-A59+4-PBA treated brains compared to MHV-A59 infected brains (Figure 4G).

Taken together, these results demonstrate the significant antiviral efficacy of 4-PBA against MHV-A59 infection in mice, effectively reducing clinical symptoms, including weight loss, hepatitis, viral titers in both the brain and the liver, as well as viral spread in the brain.

### 4-PBA modulates glial cell activation and inflammation upon MHV-A59 infection

With reduced viral spread upon 4-PBA treatment, we anticipated a difference in neuroinflammation in response to MHV-A59 infection and the activation of glial cells that are critical to neuroinflammation. To investigate this, we assessed the activation pattern of microglia and astrocytes in paraffin-embedded brain sections from both MHV-A59 and MHV-A59+4-PBA mice.

Astrocyte activation was determined in serial brain sections using immunohistochemical staining for GFAP (glial fibrillary acidic protein), the most widely used marker of reactive astrocytes (Figure 5 A, B). The quantification of GFAP staining intensity revealed significantly lower immunoreactivity in MHV-A59+4-PBA mice, consistent with lower viral load compared to MHV-A59 infected mice (Figure 5 E). Immunofluorescence analysis further confirmed there was an overall reduction in GFAP staining/ astrocyte activation in MHV-A59+4-PBA mice (Figure 5 C,D).

**Figure 5.**
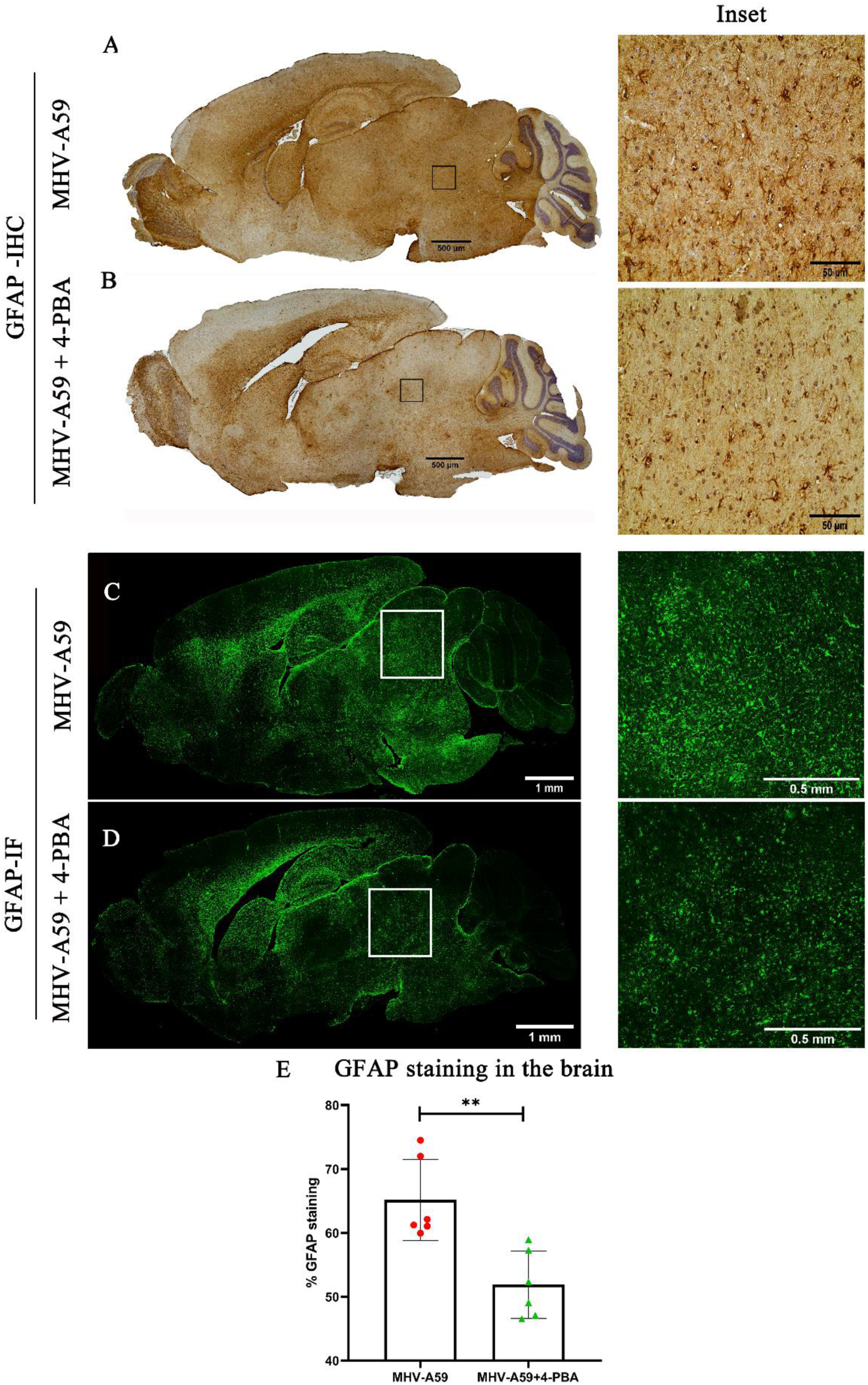
4-PBA treatment reduces astrocyte activation upon MHV-A59 infection in the mouse brain. (A,B) On day 5p.i , serial mid-sagittal brain sections (5 μm thick) from MHV-A59(A) and MHV-A59+4-PBA(B) treated mice were immunohistochemically(IHC) stained with anti-GFAP antibody. The boxed areas are shown at higher magnification alongside the corresponding brain midsagittal sections. (C, D) Photomicrographs of Serial mid-sagittal brain sections (5 μm thick) from MHV-A59(C) and MHV-A59+4-PBA(D) mice stained by immunofluorescence with anti-GFAP antibody. The boxed areas are shown at higher magnification alongside the corresponding brain midsagittal sections (E) Shows quantification of GFAP antigen staining by IHC in the brain. Results were expressed as mean ± SEM from 2 independent biological experiments (n=6 per group). *Asterisk represents statistical significance calculated using unpaired Student’s t-test and Welch correction, p<0.05 was considered significant, **p<0.01.

Microglia/macrophage activation was immunohistochemically assessed in serial brain sections with a pan-macrophage marker, anti-Iba1 (Ionized binding adaptor protein-1), confirming Iba1+ microglia/macrophage in the brain parenchyma of both MHV-A59 and MHV-A59+4-PBA mice (Figure 6 A, B). The quantification of staining intensity revealed a significant increase in the activation of Iba1+ microglia/macrophages in MHV- A59+4-PBA mice as compared with MHV-A59-infected mice (Figure 6E). Immunofluorescence analysis with anti-Iba1 further confirmed increased Iba1 staining throughout the brain of MHV-A59+4-PBA mice when compared with MHV-A59-infected mice that did not receive 4-PBA treatment (Figure 6C, D).

**Figure 6.**
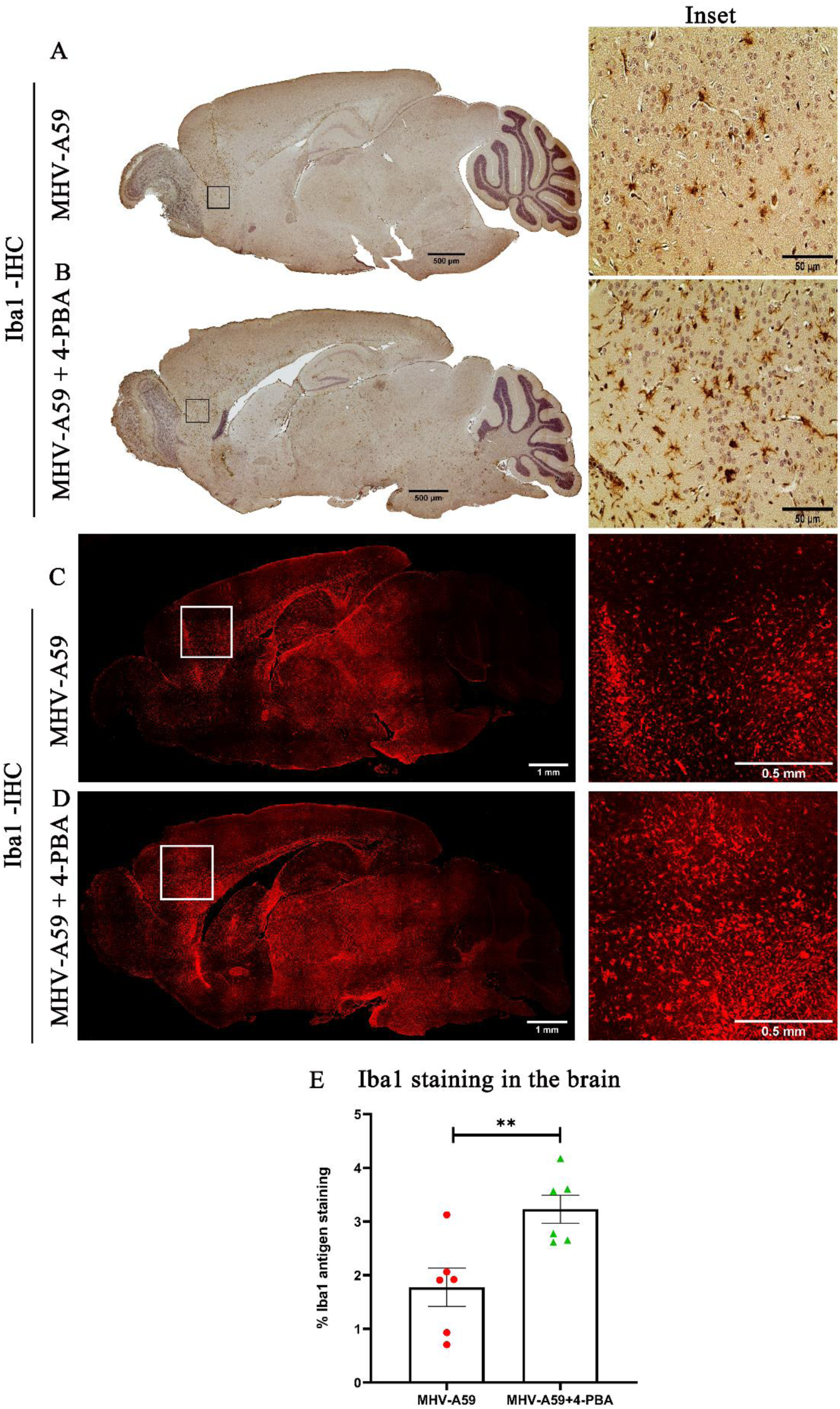
Iba1^+^ Microglia/Macrophage activation in response to MHV-A59 infection is enhanced upon 4-PBA treatment. (A,B) On day 5p.i , serial mid-sagittal brain sections (5 μm thick) from MHV-A59(A) and MHV-A59+4-PBA(B) mice were immunohistochemically(IHC) stained with Iba1 antibody. The boxed areas are shown at higher magnification alongside the corresponding brain midsagittal sections. (C, D) Photomicrographs of Serial mid-sagittal brain sections (5 μm thick) from MHV-A59(C) and MHV-A59+4-PBA(D) mice stained by immunofluorescence with anti-Iba1 antibody. The boxed areas are shown at higher magnification alongside the corresponding brain midsagittal sections (E) Shows quantification of Iba1 antigen staining by IHC in the brain. Results were expressed as mean ± SEM from 2 independent biological experiments (n=6 per group). *Asterisk represents statistical significance calculated using unpaired Student’s t-test and Welch correction, p<0.05 was considered significant. **p<0.01.

Increased microglia/macrophage activation despite reduced viral load in MHV-A59+4-PBA-treated mice was unexpected and suggested the need to study the inflammatory milieu in the CNS at this acute stage of infection when neuroinflammation is at its peak. Analyzing mRNA levels of selected inflammatory cytokines, we observed that, in line with heightened microglia/macrophage activation, mRNA expression levels of TNF-α, IL-1β, IFN-γ, and TGF-β were significantly upregulated in infected brains upon 4-PBA treatment. By contrast, IL-6 was downregulated in MHV-A59+4-PBA mouse brains when compared to that of MHV-A59-infected mice, while IL- 10 did not show any significant difference (Figure 7 A-F).

**Figure 7.**
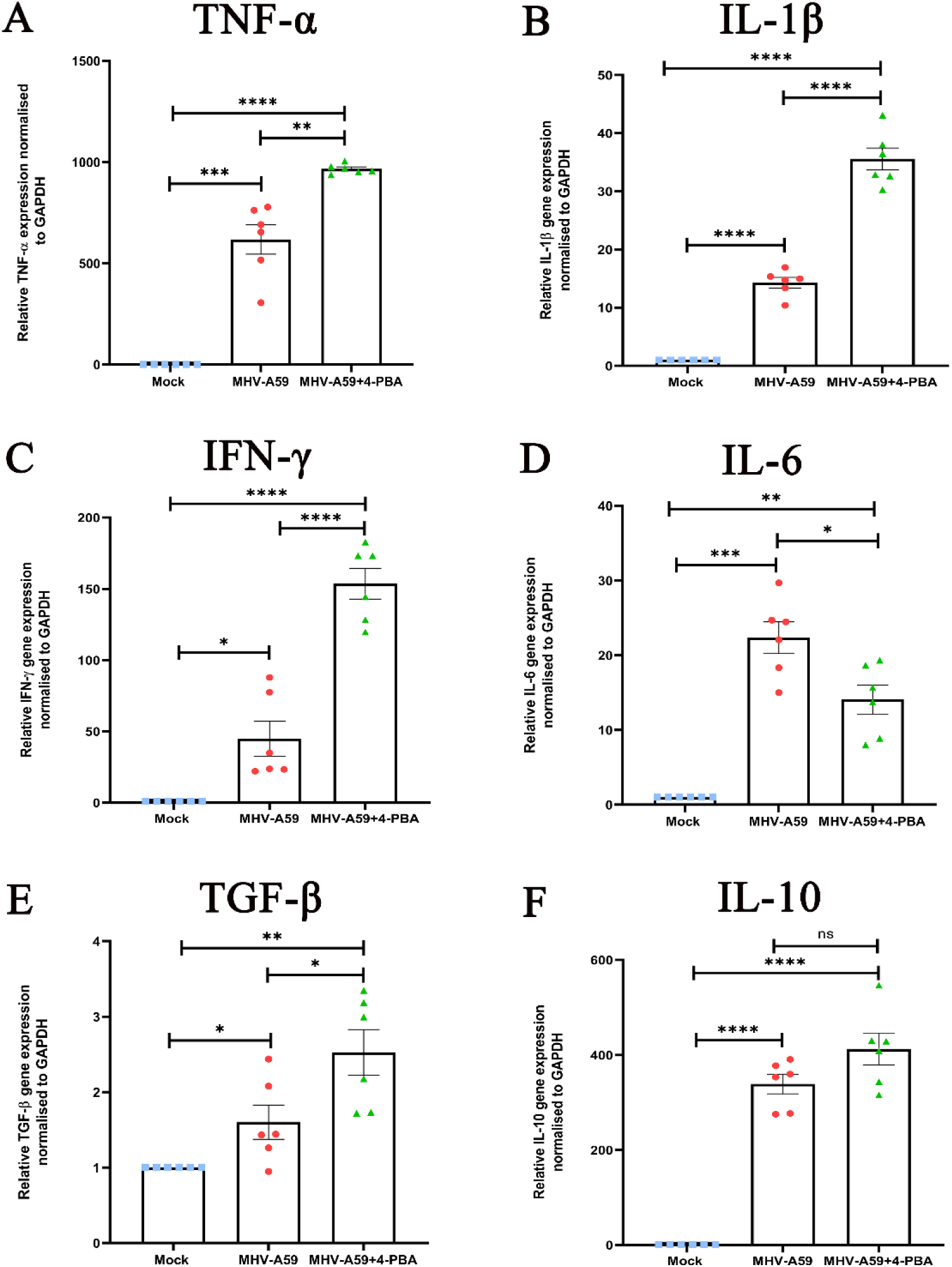
Differential mRNA expression levels of selected inflammatory cytokines in brains of infected and 4-PBA-treated mice during the acute stage of infection. RNA extracted from individual brain tissues of MHV-A59 infected and 4-PBA treated mice at Day 5p.i. was analysed for mRNA levels of the indicated cytokines by Quantitative PCR . Results were expressed as mean ± SEM (n=6 per group). Asterisk (*) represents statistical significance calculated using unpaired Student’s t-test and Welch correction, p<0.05 was considered significant. (*P<0.05, **P<0.01, ***P<0.001,**** P<0.0001).

### 4-PBA treatment mitigates the downregulation of Cx43 and ERp29 in the brain upon MHV-A59 infection

Further, we studied whether 4-PBA treatment was able to modulate Cx43 expression in the infected CNS. Immunofluorescence analysis using anti-Cx43 (Figure 8A,E; red) and anti-viral Nucleocapsid (Figure 8 B,-F, green) antibodies revealed that MHV-A59-infected mice showed a loss of Cx43 expression in infected cells of the brain (Figure 8 A-D, indicated by arrows). By contrast, MHV-A59+4-PBA-treated mice showed punctate Cx43 expression even in infected cells (Figure 8 E-H, indicated by arrows). MHV-A59+4-PBA mice showed significantly higher Cx43 gene expression compared to MHV-A59 infected mice (Figure 8 I). Immunoblot analysis confirmed the downregulation of Cx43 at the protein level in MHV-A59-infected mice, which was mitigated by 4-PBA treatment. Cx43 protein levels were downregulated in MHV-A59-infected mice as compared to mock-infected mice, while MHV-A59+4-PBA-treated mice showed levels of Cx43 protein expression comparable to mock-infected control mice. In addition, Cx43 protein levels in MHV-A59+4-PBA-treated mice were higher than MHV-A59 MHV-A59-infected mice without 4-PBA treatment (Figure 8 J, K). Thus, 4-PBA treatment could alleviate the MHV-A59-mediated downregulation of Cx43 in the brain at the acute stage of infection.

**Figure 8.**
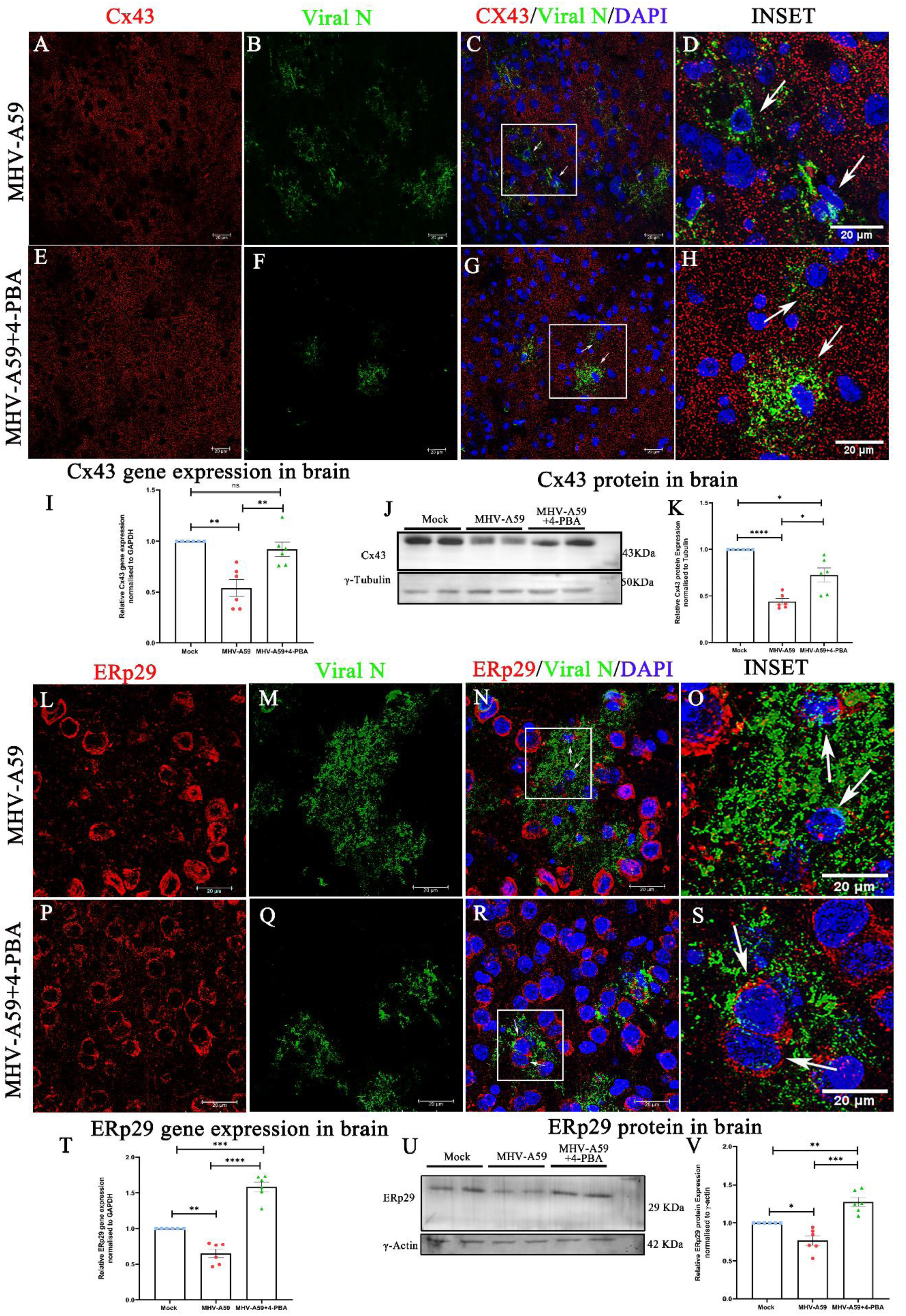
4-PBA treatment maintains Cx43 and ERp29 expression in the ERp29 in the brain upon MHV-A59 infection. A–H, At day 5p.i expression of Cx43 in the brain of MHV-A59 infected (A-D) and MHV-A59 infected+4-PBA treated (E-H) mice was analysed by double immunolabeling with anti-Cx43 (red) and anti-nucleocapsid (N) antibodies (green) followed by confocal immunofluorescence microscopy. (C-D and G-H) Arrows indicate Cx43 labelling in infected cells. I, Scatter plot of differential Cx43 mRNA in MHV-A59–infected and MHV-A59 infected+4-PBA mouse brain measured by qPCR. J and K, representative immunoblot and corresponding scatter plot of Cx43 protein expression of MHV-A59 infected and MHV-A59 infected+4-PBA treated mouse brains. L– S, Expression of ERp29 in the brain of in MHV-A59 infected (L-O) and MHV-A59 infected+4-PBA treated(P-S) mouse analysed by double immunolabeling with anti-ERp29 (red) and anti-nucleocapsid (N) antibodies (green) followed by confocal immunofluorescence microscopy. (N-O and R-S) Arrows indicate ERp29 labelling in infected cells. T, Scatter plot of differential ERp29 mRNA in Mock, MHV-A59–infected and MHV-A59 infected+4-PBA mouse brains measured by qPCR. U and V, representative immunoblot and corresponding scatter plot of Cx43 protein in Mock, MHV-A59 infected and MHV-A59 infected+4-PBA treated mouse brains. Results were expressed as mean ± SEM. Results were expressed as mean ± SEM from 2 independent biological experiments (n=6 per group). *Asterisk represents statistical significance calculated using unpaired Student’s t-test and Welch correction, p<0.05 was considered significant. **P<0.01, ***P<0.001,**** P<0.0001.

We next investigated the status of ERp29 expression in the infected CNS. MHV-A59 infection in the brain showed a loss of ERp29 in infected cells (Figure 8 L-O, indicated by arrows) as determined by immunofluorescence using anti-ERp29 (Figure 8 L, P red) and anti-viral nucleocapsid antibodies (Figure 8 M, Q, green). By contrast, immunofluorescence analysis of MHV-A59+4-PBA brains (Figure 8 P-S, indicated by arrows) demonstrated that 4-PBA treatment maintained ERp29 expression even in infected cells. This was further confirmed by qPCR analysis of ERp29 mRNA levels, which showed that ERp29 was significantly downregulated upon MHV-A59 infection. However, 4-PBA treatment in MHV-A59+4-PBA mice showed increased ERp29 that was significantly upregulated when compared to even mock-infected mice (Figure 8 T). A similar trend in expression was observed at the protein level (Figure 8 U, V).

Overall, these results demonstrate that 4-PBA treatment maintains Cx43 expression levels *in vivo*, even in MHV- A59 infected cells in the CNS.MHV-A59 infection in the CNS not only causes downregulation of Cx43 expression but also downregulates it molecular chaperone ERp29. 4-PBA treatment preserves the expression of ERp29 even in MHV-A59 infected cells in the CNS.

### 4-PBA treatment also stabilizes the expression of Cx47 in the infected CNS

Additionally we anaylsed the expression of the oligodendorcytic coupling partner of Cx43, Cx47.We analyzed Cx47 expression in the brains of MHV-A59 and MHV-A59+4-PBA mice by immunofluorescence microscopy, using anti-Cx47 (Figure 9 A,E, red) and anti-viral nucleocapsid protein antibodies (Figure 9 B,F, green). Consistent with previous reports, Cx47 plaques were disrupted in infected cells in the brain of MHV-A59 mice (Figure 9 A-D, indicated by arrows). 4-PBA treatment, which rescued Cx43 expression, also rescued Cx47 expression (Figure 9 E-H, indicated by arrows) since intact Cx47 plaques were observed in infected cells in MHV- A59+4-PBA mice brains at the acute stage of infection.

**Figure 9.**
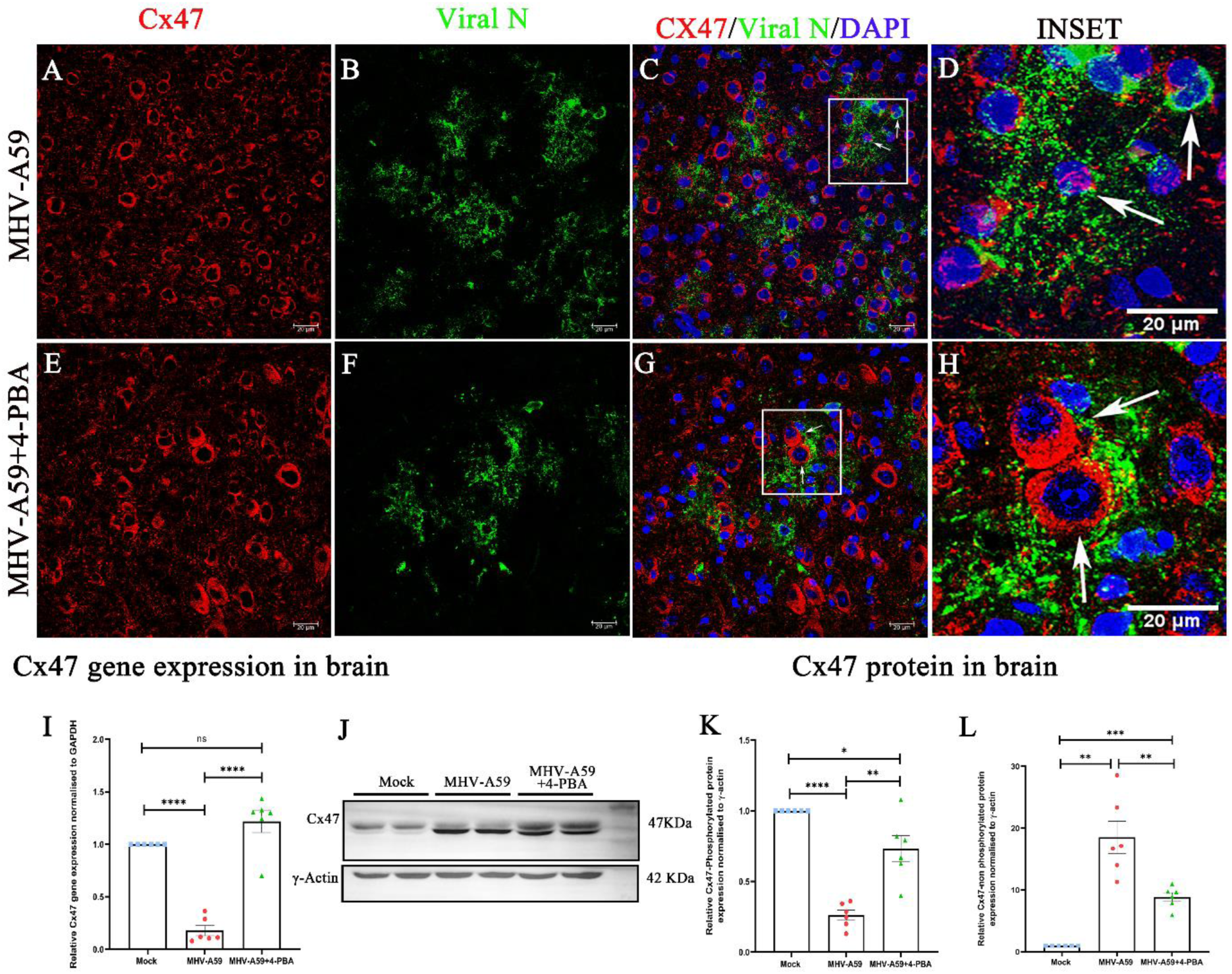
4-PBA treatment stabilises Cx47 expression in the brain upon MHV-A59 infection. A–H, Expression of Cx47 in the brain of in MHV-A59 infected (A-D) and MHV-A59 infected+4-PBA treated(E-H) mice analysed by double immunolabeling with anti-Cx47 (red) and anti-nucleocapsid (N) antibodies (green) followed by confocal immunofluorescence microscopy. (C-D and G-H)Arrows indicate Cx47 labelling in infected cells.(I) Scatter plot of differential Cx47 mRNA expression in MHV-A59–infected and MHV-A59 infected+4-PBA mouse brains measured by Quantitative PCR. (J,K and L) representative immunoblot and corresponding scatter plot of (K)Upper band(Phosphorylated) and (L) lower band(non-phosphorylated) Cx47 protein of MHV-A59 infected and MHV-A59 infected+4-PBA treated mouse brains. Results were expressed as mean ± SEM from 2 independent biological experiments (n= 6 per group). *Asterisk represents statistical significance calculated using unpaired Student’s t-test and Welch correction, p<0.05 was considered significant. *P<0.05, **p<0.01,**** P<0.0001.

Further analyzing the differential gene expression of Cx47 in MHV-A59 and MHV-A59+4-PBA mice revealed that Cx47 gene expression is significantly downregulated upon MHV-A59 infection and that this downregulation in was rescued by 4-PBA treatment. No difference in Cx47 gene expression was observed between mock-infected and MHV-A59+4-PBA-infected mice. (Figure 9 I)

Immunoblot analysis demonstrated, Cx47 was resolved as a closely migrating doublet of bands, which were quantified separately (Figure 9 J-L), since they may reflect differential Cx47 phosphorylation [39, 40]. The upper Cx47 band decreased upon infection with MHV-A59, while it did not change much in MHV-A59+4-PBA mice (Figure 9 J, K). By contrast, the lower band increased in MHV-A59-infected mice as well as MHV-A59+4-PBA mice when compared to mock-infected mice, where upregulation was significantly higher in MHV-A59 mice compared to MHV-A59+4-PBA mice (Figure 9 J, L). Thus, 4-PBA treatment also stabilized the levels of Cx47.

### 4-PBA treatment protects mice from virus-induced inflammatory demyelination at the chronic phase (day 30 p.i.)

We then determined whether 4-PBA treatment could also influence the chronic stage of demyelination in MHV- A59 infection. To study this, we treated mice with 200 mg/kg body weight/day 4-PBA until day 10 p.i. with parallel MHV-A59 infected mice as described earlier. The mice were sacrificed at day 30 p.i. and tissues were harvested to investigate demyelination at this stage.

Luxol Fast Blue (LFB) staining performed on spinal cord sections of MHV-A59 and MHV-A59+4-PBA treated mice, revealed significantly lower demyelination in 4-PBA treated mice (Figure 10 B) compared to MHV-A59 infected mice (Figure 10 A). The demyelination quantification revealed that the percentage of demyelination in MHV-A59+4-PBA treated mice was significantly lower compared to MHV-A59 infected mice (Figure 10 C). Since microglia/macrophage-mediated myelin stripping is the primary mechanism of demyelination in the MHV-induced chronic demyelination model, we next investigated the microglia/macrophage activation in spinal cords of these mice [36]. The demyelinating plaques showed a distinct accumulation of Iba1 ^+^ microglia/macrophages in serial spinal cord sections of MHV-A59 (Figure 10 D) infected mice while the Iba1 staining was lower in the spinal cords of MHV-A59+4-PBA treated mice (Figure 10 E). Quantification of the Iba1 antigen staining by immunohistochemistry shows significantly reduced Iba1 staining in the spinal cords of MHV-A59+4PBA treated mice compared to MHV-A59 infected mice (Figure 10 F). RNA from day 30 p.i. spinal cords was subjected to qPCR analysis to assess microglial phagocytic marker transcript levels of TREM2 and CD206. There was reduced mRNA expression of TREM2 (Figure 10 G) and CD206 (Figure 10 H) in spinal cords of MHV-A59+4-PBA treated mice compared to MHV-A59 infected mice. Thus, the results imply that 4-PBA treatment effectively protected mice against MHV-A59-mediated chronic stage demyelination.

**Figure 10.**
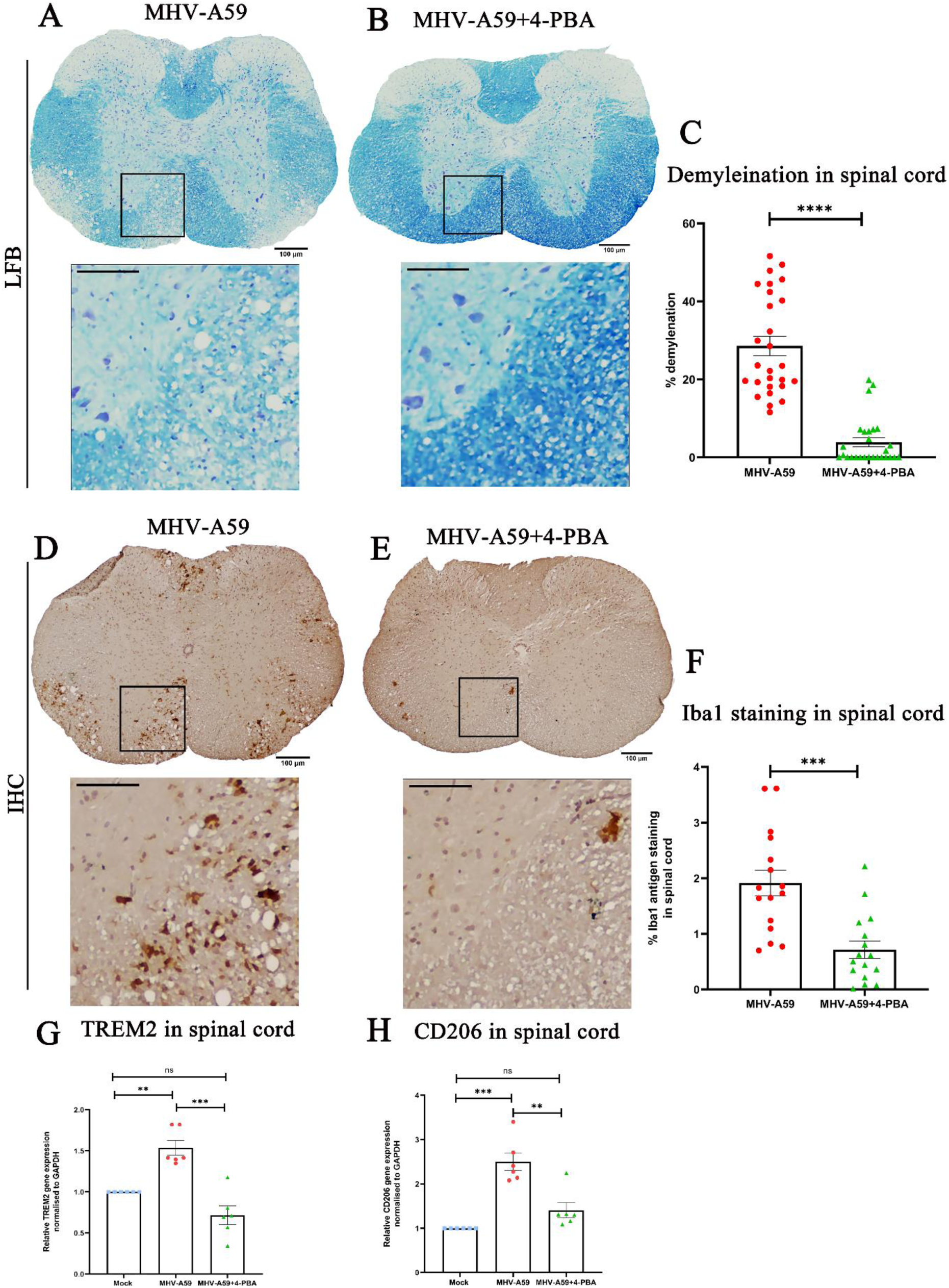
4-PBA treatment mitigates MHV-A59 mediated chronic stage demyelination. A) On day 30 p.i, cross-sections of MHV-A59 infected (A and D) and MHV-A59 infected+4-PBA treated (B and E) mouse spinal cords were analysed for demyelination by LFB and the presence of microglia/macrophages in the demyelinating lesions by Iba1 by immunohistochemistry. Black-boxed areas represent higher magnification below the corresponding spinal cord cross-sections. Scale bar 100μm, 50μm. Quantification of (C)percent area of demyelination and (F) Iba1 staining intensity. The relative transcript abundance of (G) TREM-2 and (H) CD206 was analysed in the infected spinal cords using qPCR and compared between MHV-A59 infected and MHV-A59 infected+4-PBA treated on day 30 p.i Results were expressed as mean ± SEM from 2 independent biological experiments (n=6 per group). *Asterisk represents statistical significance calculated using unpaired Student’s t-test and Welch correction; p<0.05 was considered significant. *p<0.05, ***p<0.001, ****p<0.0001, **** P<0.0001.

### Demyelinating plaque of human MS patient shows reduced staining intensity for Cx43 and ERp29

As mentioned earlier, previous studies in MS have highlighted the alteration of Cx43 in demyelinating lesions of MS patients[8, 9, 11]. Since our *in vitro[26]* and current *in vivo* studies have pointed out a concurrent loss of Cx43 and ERp29.We further validated the clinical relevance of Cx43 and ERp29 by studying archival autopsied formalin fixed paraffin embedded brain tissue sections from 1 MS case (21-year-old female) and 1 age-matched normal control (21-year-old male). Three separate plaques from the frontal and occipital lobes of the brain of the same MS brain tissue were studied. The aim was to understand the status of Cx43 and ERp29 expression in the demyelinating plaques of MS brain tissue in comparison with age-matched control. The demyelinating plaque was demarcated and characterized using loss of LFB staining in the white matter region indicative of myelin loss (Supplementary Figure 2 B). Inflammatory microglia/macrophages surrounding the plaque are characterized by the Iba1 immunohistochemical staining (Supplementary Figure 2 C). Standard control LFB staining indicated well-preserved myelin (Supplementary Figure 2 A).

Cx43 staining in the normal control brain section was seen as fluffy neuropil labelling in grey matter with scarce labelling of glia in subcortical white matter (Supplementary Figure 3 A). Demyelinating plaque in the case of MS shows a marked reduction in Cx43 staining intensity in the centre of the demyelinating plaque (Supplementary Figure 3 *, B & C) with increased staining intensity along the periphery (Supplementary Figure 3 #, B & D).

ERp29 staining in age-matched control shows the labelling of neuronal cytoplasm in grey matter and glial nuclei in white matter (Supplementary Figure 4A). Demyelinating plaque in the case of MS shows marked loss of ERp29 staining intensity in the centre of the demyelinating plaque (Supplementary Figure 4 *, B & C) with higher staining intensity within cytoplasm and nuclei along the periphery of the plaque (Supplementary Figure 4 #, B&D).

The case of MS presented here thus indicates the reduced staining intensity of both Cx43 and ERp29 compared to normal healthy control.

## 4. Discussion

This study uncovers four pivotal and previously unreported findings: 4-PBA effectively restricts the *in vivo* spread and infectivity of MHV-A59 at acute stage; acute-MHV-A59 infection in the CNS downregulates the expression of ERp29, a Cx43 molecular chaperone; 4-PBA treatment was able to maintain expression of ERp29, Cx43 and Cx47 even in infected cells at an acute-stage in the MHV-A59-infected CNS, 4-PBA significantly reduces the MHV-A59 infection-mediated virus-induced neuroinflammatory demyelination at the chronic-stage of infection.

In recent years, efforts have been focused on identifying mechanisms leading to chronic demyelination and neurodegeneration in MS to develop novel therapeutics. One promising avenue involves understanding the dynamics of gap junction proteins Cx43 and Cx47 as potential therapeutic targets for MS [9, 12]. In this context, the current study explores the *in vivo* modulation of Cx43 expression to elucidate its role in myelin pathology in a pre-clinical viral mouse model of MS. To modulate Cx43 expression in the mouse CNS upon MHV-A59 infection, we used 4-PBA.

Our study is the first to demonstrate the antiviral efficacy of 4-PBA against a β-coronavirus *in vivo* in the mouse CNS. The novelty of this study lies in demonstrating the anti-viral potential of 4-PBA against murine β- coronavirus MHV-A59 through a distinct mechanism involving ERp29 and Cx43. 4-PBA has demonstrated anti- viral effects against the Japanese Encephalitis Virus, Hepatitis C virus, and Herpes simplex type 1 virus through mechanisms such as ER stress inhibition, histone deacetylase inhibition and CREB3 downregulation, respectively[41–43]; its ability to stabilise the expression of ERp29 and gap junction proteins Cx43 and Cx47 *in vivo* in the mouse CNS represents a unique and previously unreported finding.

In a previous *in vitro* study, we demonstrated the ability of 4-PBA to rescue Cx43 expression and trafficking and restrict MHV-A59 infectivity[26]. Following this, a recent study in La Crosse Virus infection also harnessed this potential of 4-PBA, demonstrating induction of Cx43 expression and reduced viral load at early stages of infection; however, the mechanism driving 4-PBA-mediated induction of Cx43 expression remains unexplored[44]. Our study addresses this gap by demonstrating that the 4-PBA-mediated improvement in Cx43 expression correlated with enhanced ERp29 expression, which is a well-known molecular chaperone of Cx43. Thus suggesting that the improved Cx43 expression in 4-PBA-treated mice may be likely attributed to the improved ERp29 expression in these mice.

Differential glial cell activation in MHV-A59+4-PBA and MHV-A59 infected mouse brains reveals an interesting contrast. While 4-PBA-treated mice exhibit reduced astrocyte activation correlating with lower viral loads, they display heightened microglial activation. In models of hypoxia/reoxygenation injury, spinal cord injury, and Alzheimer’s disease, the deletion of Cx43 has been associated with reduced microglial activation [45–47] . Furthermore, in the model of spared nerve injury-induced neuropathic pain, astrocyte Cx43 played a significant role in regulating microglial activation [48].This suggests that increased Cx43 expression upon 4-PBA treatment likely enhances microglial activation, leading to increased secretion of inflammatory cytokines mounting a robust inflammatory response that aids viral clearance.

Cx43 plays a multifaceted role in inflammation, with studies linking Cx43 hemichannels with increased inflammation in models of acute lung injury, corneal epithelial wounding, diabetes and diabetic retinopathy, Parkinson’s and Alzheimer’s disease [45, 49]. However, in contrast to the current study, these models utilise specific Cx43 hemichannel blockers to reduce inflammation and its associated pathology. Blocking Cx43 expression using antisense oligodeoxynucleotides (asODN) to suppress Cx43 up-regulation reduces inflammation and improves functional recovery after spinal cord injury in rat models [50].

Previous studies in MHV have demonstrated that reduced immune cell infiltration in the CNS of Ifit2^-/-^ and CD4^-/-^ mice led to aggravated chronic stage demyelination[51–53]. In MHV-infected CD40L^-/-^ mice, the reduced interaction between CD4 ^+^T cells and microglia resulted in reduced microglial activation and CD4^+^T cell infiltration in the CNS, causing failure of glial cells to restore homeostasis and leading to persistently activated microglia and macrophages thereby worsening chronic demyelination pathology[54]. Thus highlighting the need for an effective transition from innate to adaptive immunity and ensuring the restoration of CNS homeostasis. Cx43 is important for efficient adaptive immune response in the CNS, facilitating antigen cross-presentation at the immunological synapse critical for adaptive immune responses and effective T-cell activation[55]. The observed enhanced Iba1 activation at day 5 p.i. and reduced chronic-stage demyelination at day 30 p.i. in 4-PBA treated mice suggests an effective resolution of inflammation following acute-stage viral clearance. Detailed studies on immune cell infiltration across various stages of infection can further elucidate how 4-PBA-mediated Cx43 expression influences adaptive immune response and viral clearance. These findings underscore the complexity of Cx43’s role in neuroinflammation and its potential as a therapeutic target, balancing the need for inflammation to clear pathogens with the necessity of resolving inflammation to prevent chronic CNS damage.

Another mechanism by which 4-PBA can reduce MHV-A59-induced chronic-stage demyelination and axonal loss is by stabilising the expression of the oligodendrocytic Cx47, improving Cx43-Cx47 mediated GJIC, which is important for proper myelination in humans as well as rodents[20]. In patients with Balo’s disease, a concentric variant of MS, astrocytic Cx43 and oligodendrocytic Cx43/Cx47 expression was diminished in both demyelinated and preserved myelin layers. NMO and MS patient samples showed preferential loss of astrocytic Cx43 in actively demyelinating and chronic active lesions, where heterotypic Cx43/Cx47 astrocyte-oligodendrocyte gap junctions were lost [8–10]. Such changes were also observed in important animal models of MS, EAE, and MHV.

In myelin oligodendrocyte glycoprotein (MOG) and myelin basic protein (MBP)-induced experimental autoimmune encephalomyelitis (EAE), levels of astrocytic Cx43 and Cx47 on mature oligodendrocytes were significantly reduced at an early stage of EAE, while Cx43 expression was increased at late EAE stages[56]. In this model, treatment with a pharmacological blocker of Cx43 hemichannels in mice, INI-0602, attenuated acute and chronic EAE. [57]. However, it should be considered with caution that these models typically lack a pathogen or infectious agent, where inflammation is crucial for pathogen clearance. Thus, reducing inflammation in these non-infectious models might be therapeutic. However, in infectious models of demyelination such as MHV-A59, initial acute inflammation is necessary for viral clearance. The inflammation must, however, resolve once the virus is cleared, as unresolved inflammation in the CNS can have devastating consequences, including demyelination and impaired remyelination.

Lastly, our observations in human MS patient-derived tissues depict reduced Cx43 and ERp29 staining intensity in the demyelinating plaques, supports the need to study ERp29 in the context of GJIC and requires further validation in larger cohorts. This, along with the observed loss of Cx43 in MHV and EAE-induced models of MS and the loss of ERp29 in MHV-A59 induced animal model of MS, highlight the potential of Cx43 and ERp29 as important therapeutic targets for MS. To the best of our knowledge, this is the first report demonstrating ERp29 expression in the human brain and reduced staining intensity in MS plaque.

This study highlights the potential of 4-PBA to enhance ERp29 expression and stabilise the expression of gap junction proteins, mitigating chronic virus-induced demyelination in an MS model and underscores the importance of ERp29 for Cx43 expression in mice and humans. Thereby suggesting the therapeutic benefits of repurposing 4-PBA to improve GJIC in MS by regulating ERp29. Moreover, this is the first report to demonstrate 4-PBA’s ability to reduce *in vivo* β-CoV infectivity, suggesting its potential as a therapeutic agent in virus-induced CNS pathologies. The study identifies Cx43 as a novel target in designing β-CoV antivirals and addressing virus-induced demyelination.

## Acknowledgements

This work was supported by the Anusandhan National Research Foundation formerly Science and Engineering Research Board (SERB: CRG/2023/000999 and SPG/2020/000454) and internal support from IISER Kolkata. The Council for Scientific and Industrial Research (CSIR) provided fellowships to GK, MS and SK.The authors would like to acknowledge the support of the Department of Biological Sciences ,IISER Kolkata, Central Imaging Facility, and DBT-FIST imaging facility, and Ms Shreya Ghosh and Mr Ritabrata Ghosh. The authors would like to convey special thanks to Prof.Arnab Gupta, IISER Kolkata, for the Wellcome trust- Leica SP8 confocal platform and Mr Raviranjan Pandey for his help in image acquisition. The authors would like to thank Human Brain Bank, NIMHANS, Bangalore, India, for providing the human brain tissue samples. The DBT-SYMEC facility and BioInspired Pvt. ltd for providing infrastructural and equipment-related support.

