## Supplementary for "4-Phenylbutyric acid modulates Connexin 43 expression restricting murine-β-coronavirus infectivity and virus-induced demyelination"


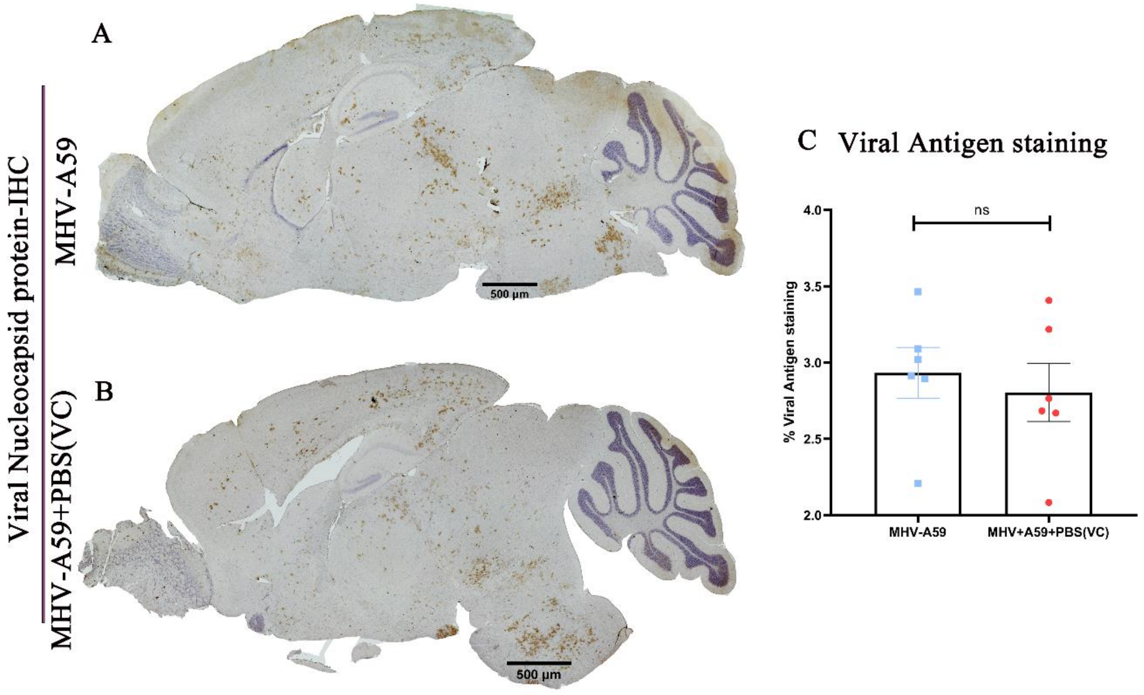


**Supplementary Figure 1:** ***Comparable Viral Antigen Staining in MHV-A59 Infected and Vehicle Control Mice***(A,B) On day 5p.i , serial mid-sagittal brain sections (5 μm thick) from MHV-A59 (A) and MHV-A59+PBS(VC) (B) sets of mice were immunohistochemically(IHC) stained with anti-N antibody.(C) Shows quantification of Viral N antigen staining by IHC thus demonstrating that no differences in viral antigen staining were observed between MHV-A59 infected and VC mice (A, B) in the brain. Results were expressed as mean ± SEM from (n=6 per group). Statistical significance was calculated using unpaired Student's t-test and Welch correction.

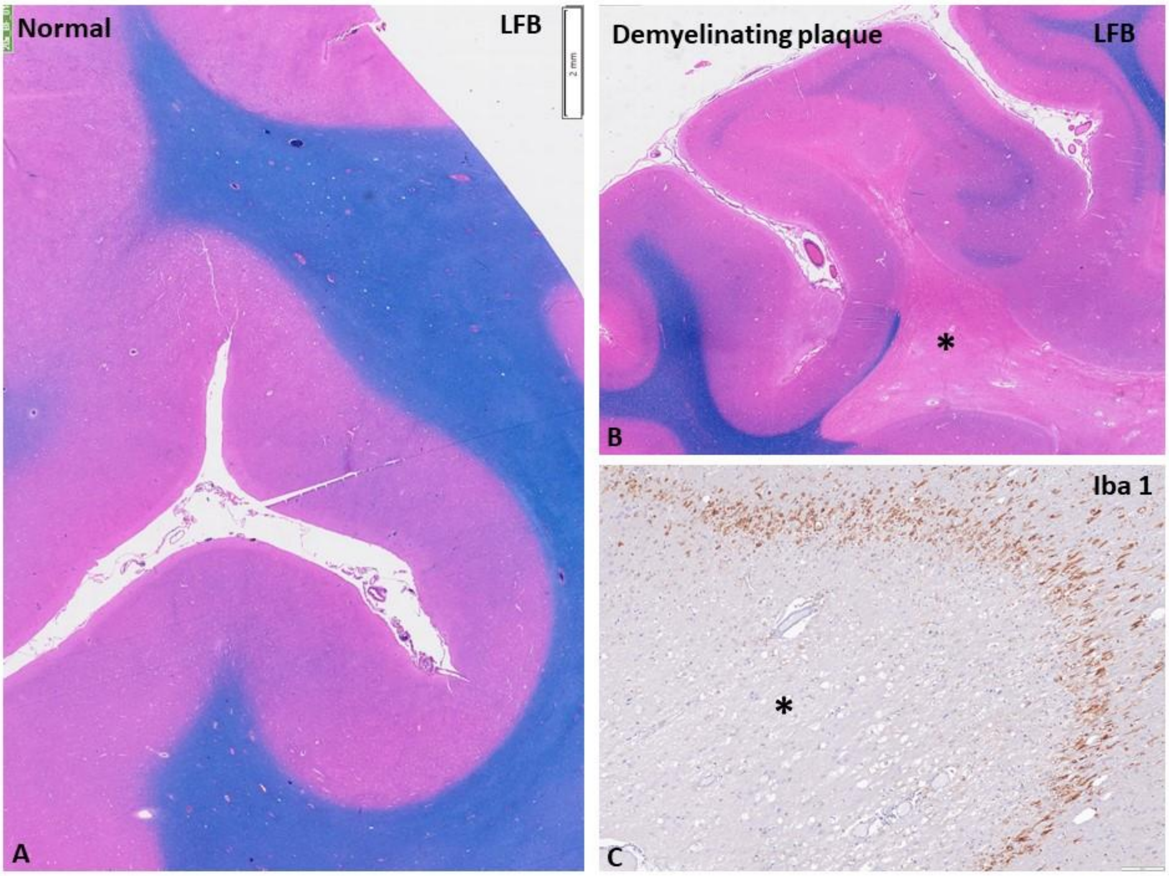


**Supplementary Figure 2: Characterisation of demyelinating plaque in white matter in the case of MS compared to age-matched normal control.**(A) Luxol fast blue stain is in normal control, showing well-preserved myelin in white matter. (B-C) Large demyelinating plaque in white matter in case of MS highlighted by LFB stain shows marked loss of expression of myelin (*, B&C) Band of macrophages are seen along the periphery of the plaque (C) highlighted by Iba1 – marker for macrophages.[magnification=scale bar]

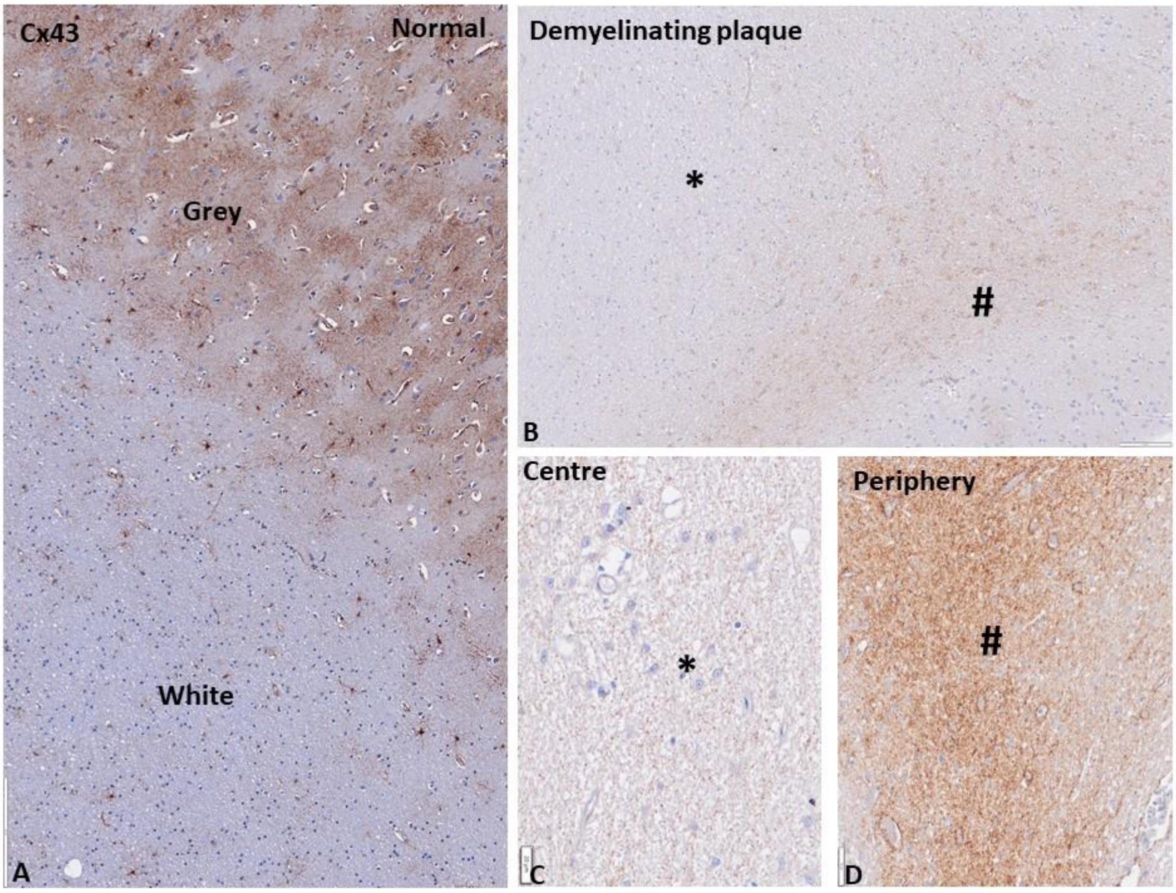


**Supplementary Figure 3: Cx43 staining as observed in the white matter of normal and MS human brain tissues.** (A) Cx43 staining seen as fluffy neuropil labelling in grey matter in normal control with scarce labelling of glia in subcortical white matter. (B-D) Demyelinating plaque in the case of MS shows a marked reduction in staining intensity of Cx43 in the centre of the demyelinated plaque (*, B & C) with increased staining intensity along the periphery of the plaque localising to macrophages/reactive glial processes (#, B&D). [magnification=scale bar]

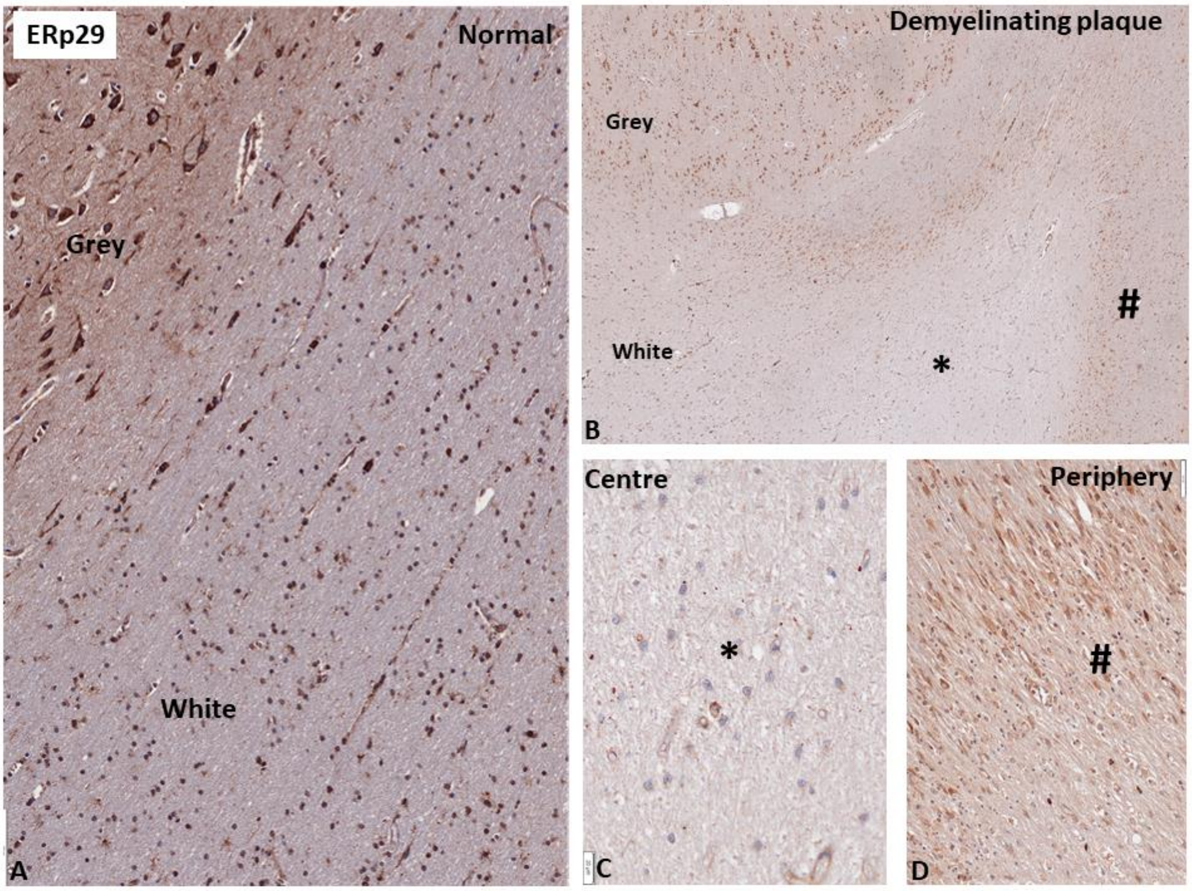


**Supplementary Figure 4: ERp29 staining as observed in the white matter of normal and MS human brain tissues.** (A) ERp29 immunolabeling in age-matched control shows the labelling of neuronal cytoplasm in grey matter and oligodendroglia and astrocytic nuclei in white matter(B-D). Demyelinating plaque in case of multiple sclerosis shows marked loss of ERp29 staining intensity in the centre of the demyelinated plaque (*, B & C) with increased staining intensity within cytoplasm and nuclei of reactive macrophages/astrogliosis along the periphery of the plaque (#, B&D). Note the preserved expression of ERP in neurons in grey matter. [magnification=scale bar]
